# Learning-Induced Effects of Practice Schedule Variability on Stimuli Discrimination Efficiency: High-Density EEG Multi-scale Analyses of Contextual Interference Effect

**DOI:** 10.1101/2024.08.29.610352

**Authors:** Alexandre Cretton, Kate Schipper, Mahmoud Hassan, Paolo Ruggeri, Jérôme Barral

## Abstract

Contextual interference (CI) enhances motor learning by practicing skill variations in a random rather than blocked order. It has been demonstrated that performing aiming distances in a random order increased electrophysiological (EEG) markers of perceptual, attentional, and working memory processes. However, only the effect of CI on these markers before training was assessed, without evaluating whether they would decrease with learning in participants trained under the random compared to the blocked condition, indicating enhanced neural efficiency. To address this, 35 participants practiced an aiming task involving three distances over nine sessions across three weeks. They were divided into two groups: one trained with distances in a random order (HCI group) and the other in a blocked order (LCI group). Electrophysiological activity was recorded for all participants in the random condition before and after the training program using a high-density EEG multiscale approach, including topographical, source estimation, and source connectivity analyses. EEG analyses revealed post-training neural dynamic differences between groups. The HCI group showed reduced and shorter P3a-like activity compared to the LCI group, while the LCI group exhibited greater occipito-temporal-frontal gamma-band synchronization. These findings suggest that random practice enhances the efficiency of perceptual and attentional processes, particularly of stimuli discrimination, compared to blocked practice.

## 1. Introduction

The contextual interference (CI) effect is a motor learning phenomenon characterized by improved learning outcomes when skill variations are practiced in a random rather than a blocked order (Battig, 1979; Shea & Morgan, 1979; Magill & Hall, 1990; Czyż et al., 2024). Under a blocked schedule (low CI), learners practice the same variation repeatedly within each block before switching to another variation in subsequent blocks. In contrast, a random schedule (high CI) intermixes all skill variations within each block. Although random practice typically results in poorer immediate performance during training, it significantly enhances performance on subsequent retention and transfer tests (Shea & Morgan, 1979; Magill & Hall, 1990; Czyż et al., 2024).

Two main hypotheses have primarily been proposed to explain the CI effect: the elaboration hypothesis (Shea & Zimny, 1983) and the forgetting-reconstruction hypothesis (Lee & Magill, 1983, 1985). According to the elaboration hypothesis, random practice enhances motor learning by facilitating inter-item comparisons during the inter-trial interval, thereby enriching memory encoding. For instance, in aiming tasks (e.g., Young et al., 1993; Ramezanzade et al., 2022; Cretton et al., 2024, Cretton et al., 2025), random schedules require maintaining multiple practiced distances in working memory to enable comparisons between them. In contrast, the forgetting-reconstruction hypothesis suggests that random practice imposes frequent switching between tasks, compelling learners to repeatedly suppress the motor plan executed on the previous trial and reconstruct a new motor response plan (e.g., direction, amplitude, force in aiming tasks) before each subsequent movement, thus strengthening long-term motor memory (Lee & Magill, 1983, 1985).

Complementarily, other research highlights that random practice enhances perceptual discrimination and attentional control, accommodating continuously changing task demands such as varying target distances (Li & Wright, 2000; Wright et al., 2016; Immink et al., 2020; Lage et al., 2021). These theoretical explanations, while emphasizing different underlying cognitive processes, collectively support the view that random practice increases overall cognitive demands. Supporting this integrated perspective, Li and Wright (2000) demonstrated that tasks requiring discrimination between auditory tones during inter-trial or pre-response periods impaired performance more significantly under random compared to blocked conditions, reflecting increased attentional resource allocation during random practice.

Neuroimaging studies also have supported the view that random practice requires greater cognitive processing than blocked practice. For instance, increased working memory demands during random practice have been inferred from greater engagement of the dorsolateral prefrontal cortex (DLPFC) compared to blocked practice (Kantak et al., 2010; Lin et al., 2011; Shewokis et al., 2017). This brain region is critically involved in managing the temporary storage and manipulation of task-relevant information within working memory (Constantinidis & Klingberg, 2016; Riley & Constantinidis, 2016).

In addition, random practice necessitates continuous updating of sensory information to accurately discriminate between changing task demands, leading to increased activation of the occipital visual cortex—a region fundamental for early visual processing (Murray et al., 2002; Murray et al., 2004). This sensory-processing requirement has also been confirmed by fMRI studies showing stronger occipital cortex activation under random practice (Pauwels et al., 2018), and further supported by reduced occipital GABA concentrations, indicative of inhibitory tuning that facilitates enhanced sensory discrimination (Chalavi et al., 2018). Electrophysiological findings further corroborate this, demonstrating more robust EEG markers of attention during random practice compared to blocked practice (Lelis Torres et al., 2017).

Independently of the specific cognitive processes considered, neural markers of contextual interference generally show greater brain activation during random practice compared to blocked practice, reflecting increased cognitive demands. Following training, however, this pattern of neural activity typically reverses. Participants who trained under random conditions exhibit lower neural activation relative to those trained under blocked conditions, indicating more efficient and effective neural processing after training (Kantak et al., 2010; Lin et al., 2010; Lin et al., 2011; Lin et al., 2013; Lin et al., 2016; Frömer et al., 2016; Shewokis et al., 2017; Chalavi et al., 2018; Immink et al., 2021).

Supporting these findings, a recent high-density EEG study (Cretton et al., 2024) provided evidence that random practice enhances neural markers of perceptual, attentional, and working memory processes compared to blocked practice. Specifically, random practice elicited stronger early perceptual responses, reflected by increased N1-like scalp topography and enhanced ventral cortical activation, suggesting greater demands for stimulus categorization and recognition. Moreover, random practice induced a prolonged P3a-like topography along with heightened anterior cingulate cortex (ACC) activity, indicative of increased attentional resources required for accurately discriminating and switching among varying aiming distances. Additionally, a pronounced P3b-like topography accompanied by elevated parietal cortical activation uniquely characterized random practice, demonstrating enhanced working memory engagement required to maintain and dynamically update spatial information about the aiming targets.

Furthermore, two neural networks exhibited significantly greater synchronization under random compared to blocked practice. The first, a ventral alpha-band network, encompassed regions such as the lingual and fusiform gyri; inferior, middle, and superior temporal gyri; temporal pole; entorhinal cortex; parahippocampal gyrus; pars opercularis; medial and lateral orbitofrontal cortex; insula; and right pars orbitalis. This network is closely associated with sustained attentional focus and vigilance. The second, a frontoparietal theta-band network, is involved in cognitive control and executive functioning. Collectively, these findings indicate that random practice conditions evoke greater activation and coordination across brain regions supporting critical cognitive functions necessary for managing the continuous task variations induced by the random condition.

However, these previous findings (Cretton et al., 2024) reflect neural markers observed only during initial task performance at baseline, specifically highlighting cognitive processes uniquely engaged in the random condition relative to the blocked condition. In the motor learning literature, a crucial distinction is drawn between immediate performance—temporary task execution influenced by transient factors such as fatigue or strategy—and actual learning, characterized by stable performance improvements measured through retention or transfer tests (Soderstrom & Bjork, 2015). While the earlier findings demonstrated increased neural activity specifically associated with performing in the random condition prior to training, the current investigation focuses explicitly on how these neural processes, initially enhanced in the random condition, are modified by extensive practice. Specifically, the present investigation examines whether participants who trained under random conditions subsequently exhibit reduced neural activity—indicative of facilitated processing—when tested again in the random condition compared to participants who trained under blocked conditions.

To address this goal, we designed a protocol involving 36 participants who completed nine training sessions distributed across three weeks. Participants practiced an aiming task involving three distinct distances under either a random (high contextual interference group, HCI group) or blocked (low contextual interference group, LCI group) practice schedule. Electrophysiological activity was recorded using high-density EEG—including scalp topography, source estimation, and connectivity analyses— one day before the start of training and again one day after completing the final training session. Critically, connectivity analyses were included to extend beyond localized neural activity measures and examine how large-scale brain networks adapt in response to extended training. While previous research has largely emphasized localized neural changes within regions such as the prefrontal and parietal cortices (for reviews, see Lage et al., 2015; Wright et al., 2016), examining connectivity allows investigation into whether contextual interference effects also modulate the coordination between these and other regions involved in cognitive and motor processes.

Importantly, all data analyzed in this study were collected from a single cohort of 36 participants who underwent nine training sessions and three test sessions, previously reported in two separate studies (Cretton et al., 2024; Cretton et al., 2025). The test sessions occurred one day before training (T1, baseline), one day after training (T2, 24 hours retention), and one week after training (T3, one week retention). During the training phase, participants exclusively practiced under their assigned conditions—either random (high contextual interference, HCI group) or blocked (low contextual interference, LCI group). During each test session, all participants performed tasks under both random and blocked conditions in a counterbalanced order. EEG recordings were conducted exclusively during these test sessions. Previous analyses from this cohort separately reported the EEG findings at baseline (T1) that were introduced earlier in the introduction, highlighting neural differences between random and blocked conditions before training (Cretton et al., 2024). Additionally, the behavioral results across training and retention sessions, described previously in the introduction, clearly demonstrated the contextual interference effect (Cretton et al., 2025). Specifically, the HCI group exhibited poorer initial performance during training but superior accuracy in both retention tests (T2 and T3) and transfer tasks compared to the LCI group. Critically, at the T2 retention test, the HCI group demonstrated a clear accuracy advantage, but only when tested under the random schedule. In the current study, we aim to extend these prior findings by specifically analyzing EEG data recorded in the random test condition at baseline (T1) and in post training in a 24-hour retention session (T2). By exclusively examining EEG data from the random condition at T2, we can directly investigate how the neural processes identified at baseline as uniquely engaged by the random condition (Cretton et al., 2024) are modulated by extensive training under random versus blocked schedules. Additionally, we target the random condition because the behavioral advantage of the HCI group was specifically observed in this condition, not in the blocked condition. Importantly, potential biases introduced by this methodological choice are explicitly addressed in the discussion section. For a full description of the experimental design and detailed behavioral analyses, readers are referred to Cretton et al. (2025). Initial EEG analyses at baseline are detailed in Cretton et al. (2024).

Based on prior EEG findings (Cretton et al., 2024), we hypothesized that extensive training under random practice conditions would lead to distinct neural adaptations compared to blocked practice. Specifically, at the ERP level, we predicted that neural markers initially enhanced during random practice—such as the N1-like, P3a-like, and P3b-like components—would exhibit a greater decrease in the HCI group compared to the LCI group when performing again under random conditions post-training. At the source level, we expected decreased activation in cortical regions previously identified as crucial for perceptual categorization, attentional switching, and working memory updating— including ventral prefrontal, parietal as well as occipito-temporal regions—in participants trained under random conditions compared to blocked conditions. Regarding functional connectivity, we hypothesized that following extensive random practice, participants in the HCI group would demonstrate reduced synchronization within previously identified neural networks: specifically, decreased connectivity in the ventral alpha-band network associated with sustained attentional focus, as well as reduced connectivity in the frontoparietal theta-band network linked to cognitive control and executive function, compared to participants in the LCI group.

## Methods

### 2.1 Participants

Between October 14, 2021, and June 2, 2022, we enrolled 36 right-handed students from the University of Lausanne. To assess the handedness, we used the Short-form Edinburgh Handedness Inventory (Veale, 2014) to obtain a notation from −100 to +100. Only participants that obtained a right-handed score of at least +80 were included in the study. These participants were quasi-randomly assigned to either the Low Contextual Interference (LCI) group or the High Contextual Interference (HCI) group. Group matching was ensured based on age, gender, and the pretest session’s mean and standard deviation of response time (reaction time + movement time) and error rates, as defined in Section 2.3. In order to calculate the number of participants for our study, we used the G-Power software (Faul et al., <u>2007</u>). To estimate the required sample size, we used the effect size reported by Hall and Magill (1995), which yielded a target of 42 participants to achieve a power of 0.80 with an effect size of f = 0.1875. We were able to recruit 38 participants, but two dropped out before the first session (prior to group assignment). Although the power analysis was based on behavioral contextual interference effects, it is accepted that EEG studies can maintain sufficient power with fewer participants, due to the increased number of trials and lower between-subject variance (Baker et al., 2021). Additionally, one participant in the HCI group was excluded due to excessive slow-wave artifacts in the EEG recording, likely caused by extreme fatigue. The final sample thus included 18 participants in the LCI group (11 females, 7 males; mean age = 21 years, SD = 3 years; age range = 18–28 years) and 17 participants in the HCI group (13 females, 4 males; mean age = 20.88 years, SD = 1.45 years; age range = 18–24 years). All participants provided written informed consent, met the eligibility criteria—including having normal or corrected-to-normal vision and no history of psychiatric or neurological disorders—and were compensated with 100 Swiss Francs (approximately 107 US dollars) for their participation. The local ethical committee approved all procedures prior to the start of the study (CER-VD: 2021-01456).

### 2.2 Task and apparatus

The task (Figure 1A) required participants to extend their left arm to move a cursor from a starting point at the bottom of the computer screen to a top target. Both the target and starting positions were represented by one-centimeter-wide and –high crosses (length in visual angle: 0°49’). Participants had to execute a straight movement along the vertical dimension of the screen and to click as close as possible to the target. In each trial, the physical distance between the starting point and target could be either 7 cm (short distance), 14 cm (mid distance), or 21 cm (long distance). The corresponding visual angles were 5°43’, 11°25’ and 17° 30’, respectively. To standardize movement execution across trials, their forearms rested on a wooden support with their elbows placed on a wooden wedge. At the beginning of each trial, the forearms formed a 90-degree angle with the upper arm, which was parallel to the body. To minimize head movement during the experiment, participants used a chin rest device. This ensured that their eyes remained fixed in the middle and 70 cm away from the screen, providing constant visual access to both the starting point and target without the need for eye movement. The position of the wooden support and the height of the seat were adjusted for each participant’s body dimensions. Additionally, participants could not initiate cursor movement prior to the start of the trial, as the cursor was automatically set and blocked to the starting position. The cursor was constrained to move along the y-axis only and could not deviate along the x-axis. To prevent visual feedback and online control of movement, neither the cursor’s displacement nor the hand displacement were visible during the pointing movements. The task was powered by the PsychoPy 2021.1.2 (Peirce et al., 2019) software and displayed on a Dell Alienware AW2521HFA computer monitor. The monitor had a refresh rate of 240Hz, a screen resolution of 2560 x 1440 pixels, and a 24.5-inch screen size. The monitor’s IPS response time of 1 ms (grey to grey) enabled smooth and precise cursor movements. Arm movements were executed using an ambidextrous computer mouse (Kova AIMO), which has dimensions of 130 mm in width, 67 mm in depth, and 40 mm in height, and weighs 100 g. The mouse features infrared tracking and was set to a resolution of 200 dpi to control the cursor. To respond participants clicked on the button with the index finger.

**Figure 1.**
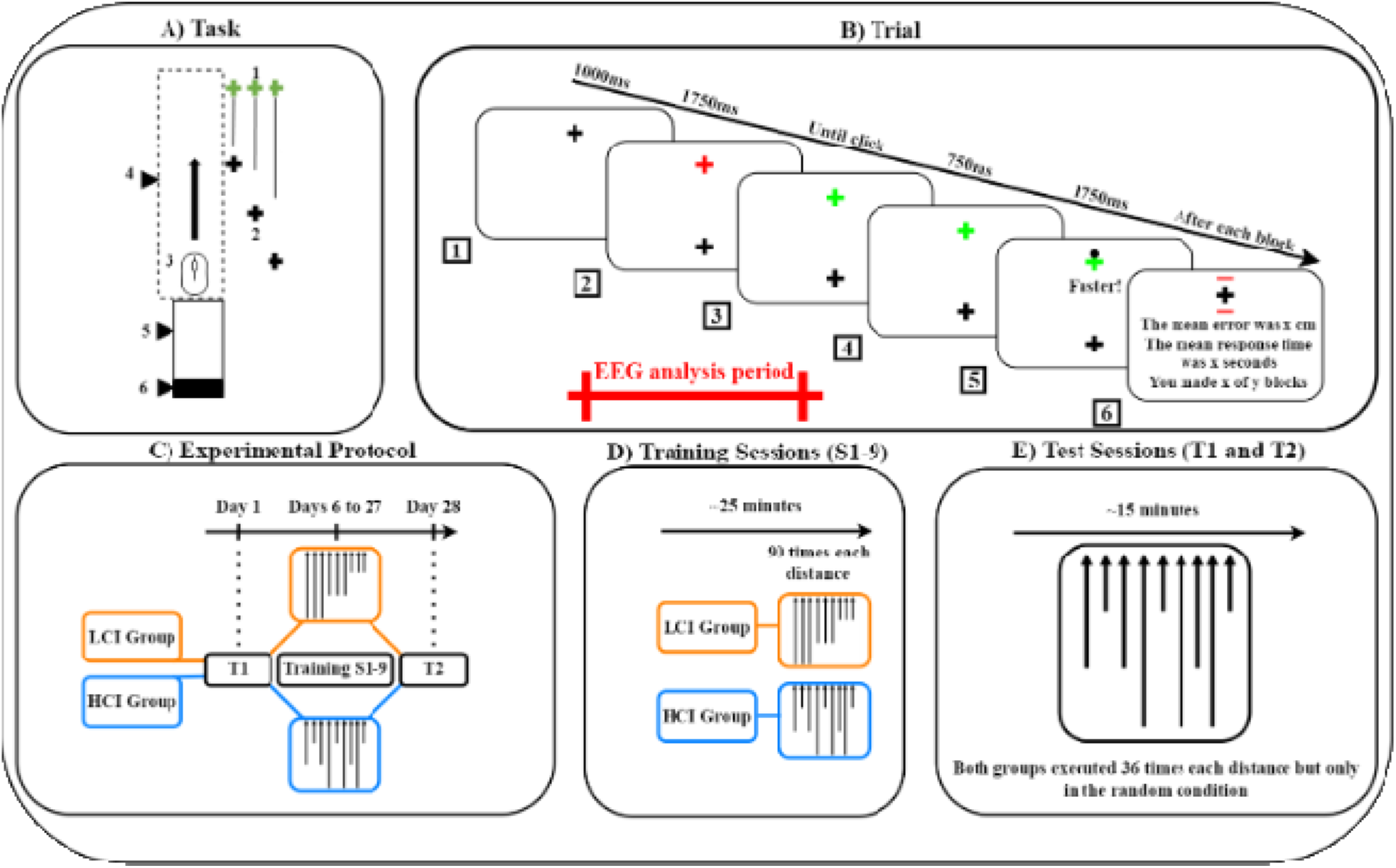
Experimental Design, Materials, and Procedures. (A) Task: The experimental setup included (1) the target, (2) starting points at short, mid, and long distances, (3) an ambidextrous computer mouse, (4) a wooden box obscuring the hand, (5) a wooden support for the arm, and (6) a wooden wedge brace for the elbow during the inter-trial interval. The computer monitor displaying the task is not depicted in this representation. (B) Trial: (1) A trial began with a fixation cross displayed for 1000ms. (2) Following this, participants prepared to move for 1750 ms when two crosses appeared simultaneously: a black starting point and a red target. (3) Once the target changed color from red to green, (4) participants swiftly moved the invisible cursor from the starting cross (located at the bottom of the screen) to the target cross (at the top) and attempted to click as close to the target as possible. (5) A 750ms pause ensued, after which feedback was shown for 1750 ms—a black dot indicating the cursor’s click position. (6) If the response time surpassed 20% of the baseline established in the familiarization trials, the black dot would not appear; instead, a prompt reading “faster” would be displayed. After each block, participants received summary feedback, which included red outlines around the target—representing the mean error in centimeters— and mean response time (in seconds) along with information about the number of completed and remaining blocks. The red bar marks the period segmented for EEG analysis, spanning from 200 ms before the red warning cross-stimulus (box 2) to 550 ms after the green imperative cross-stimulus (box 3) (described in the EEG recording and preprocessing section). (C) Experimental Protocol: On Day 1, participants completed the baseline test (T1). They then underwent 9 training sessions spanning from Day 6 to Day 27. The post-training 24-hour retention test (T2) was completed on Day 28, followed by the final test (T3) on Day 35. Participants in the HCI group trained under the random condition, while those in the LCI group trained under the blocked condition. (D) In each session, participants completed 270 trials, with 90 trials for each of the three distances in their respective condition. These trials were organized into 18 blocks, each containing 15 trials. For the HCI group, training followed the random condition schedule, where each block had 5 trials for every distance, presented in a randomized sequence. Conversely, for the LCI group, training followed the blocked condition schedule: participants trained one specific distance for 6 consecutive blocks (a total of 90 trials) before moving on to the next distance for another 6 consecutive blocks, and then finished with 6 consecutive blocks of the final distance. The order of the distance blocks in the blocked condition was counterbalanced across participants and sessions. (E) Each test session consisted of 36 trials for each distance in the random condition. Thus, all participants, irrespective of their group, performed the task in the random condition.

**Figure 2.**
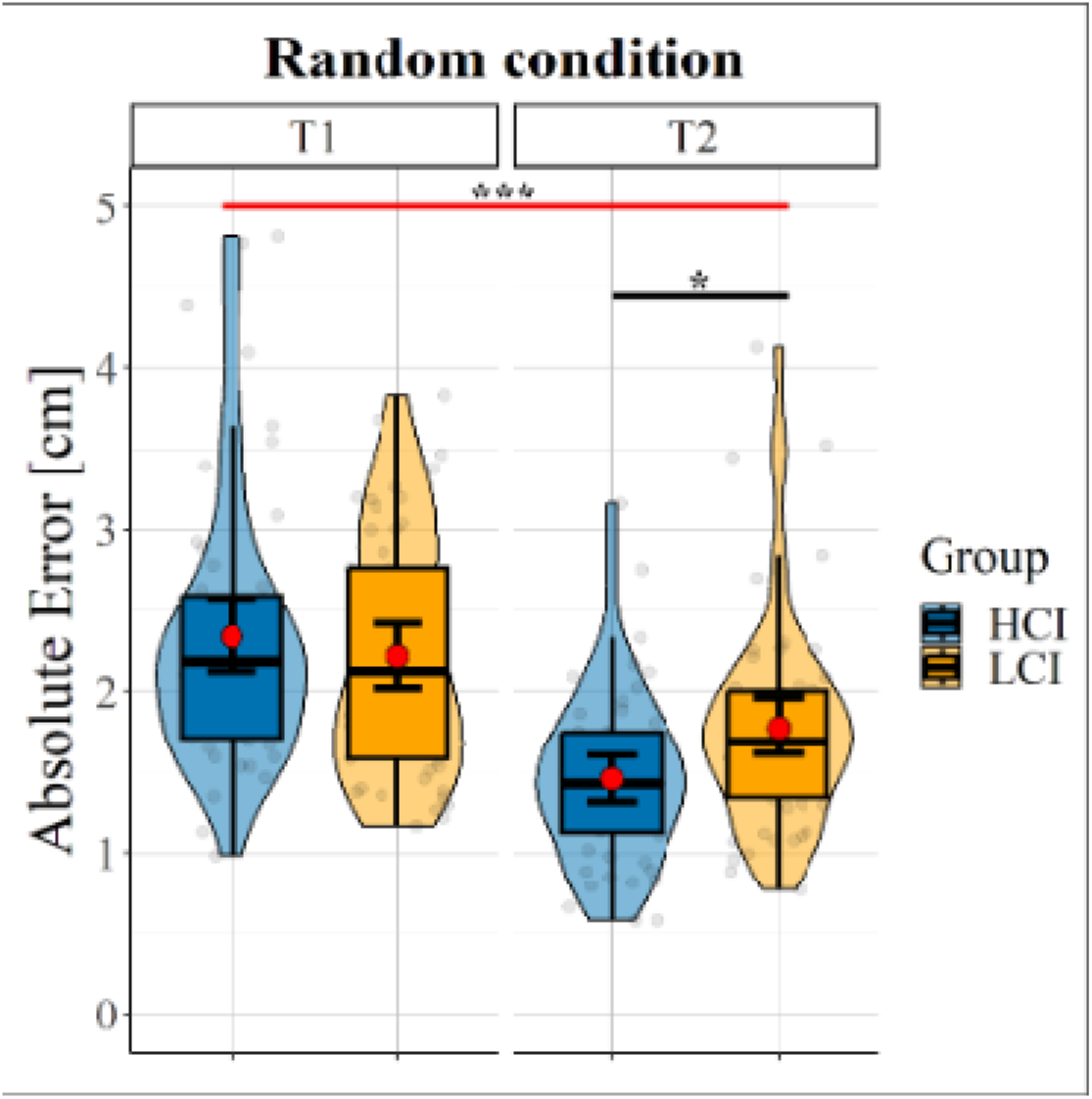
Data distribution of Absolute Error across sessions (T1, T2) for the HCI (blue) and LCI (orange) groups in the random condition. The red points represent mean values. Significant post-hoc test results (Holm-bonferroni corrected) are indicated by the red bar for the main effec s of session and by the black bar for the group × session interaction: *(p ≤ 0.05), **(p ≤ 0.01), ***(p ≤ 0.001).

### 2.3 Trial

There are two types of sessions in this experiment: test sessions and training sessions (described in the next section). The chronological sequence of one trial during the test session is depicted in Figure 1B. At the beginning of each trial, an intertrial fixation cross was displayed on the screen at the target location for 1000 ms. Then, two crosses, one indicating the starting position (in black) and the other one (in red) the target to hit were presented during 1750 ms. Participants were required to move towards the target when the color of the target cross changed from red to green. The movement execution period remained open until the participant clicked the mouse button in the vicinity of the target. After 750 ms, immediate feedback – a black dot – appeared on the screen at the location of the click. If the total trial time (i.e., reaction time + movement time) exceeded 20% of the average time computed during familiarization (performed at the beginning of each task), the black dot feedback was replaced by a message “faster” at the center of the screen. The black dot or the message were displayed for 1750 ms. Finally, at the end of each block, feedback displaying the mean values of error and response time was presented along with the number of completed and remaining blocks. Simultaneously, a graphical representation of the mean error drawn around the target was displayed (red lines on the last panel of Figure 1E), allowing participants to self-evaluate their performance. At this moment, they could choose to take a break or move to the next block by pressing the space bar.

During the training sessions, the trial’s structure was similar, with the following exceptions: the intertrial fixation cross duration was randomized between 300 and 700 ms, the movement preparation period was randomized between 700 to 1700 ms, and the feedback period was displayed for 750 ms. These changes shortened the duration of the experimental sessions, reducing the burden on the participants.

### 2.4 Experimental protocol and instructions

The experimental protocol comprised nine training sessions and two test sessions over a 28-day period (Figure 1C). All sessions took place at the Laboratory of Experimental Research on Behavior within the University of Lausanne’s facilities. During the test sessions, both EEG and performance were recorded, whereas only performance was recorded during the training sessions. On Day 1, a pretest session (T1) was conducted to establish a baseline for each participant’s task-related cortical electrophysiological activity and performance. From Days 6 to 27, participants completed nine training sessions (S1-9). On Day 28, a post-training 24-hour retention session (T2) evaluated the participants’ task-related cortical electrophysiological activity and performance following the training program. The pretest and 24-hour retention sessions were identical. At T1, participants received written instructions on how to perform the task and the experiment’s goal. They were not informed of the experimental conditions and were only instructed to improve their task performance. Although participants were aware that the trial times were recorded, the focus was on improving accuracy. Test sessions were conducted at the same time and on the same day of the week for each participant at both T1 and T2, maintaining a consistent schedule per participant. In contrast, participants could complete their training sessions anytime between 8:00 a.m. and 5:00 p.m., Monday through Friday, without the need for an appointment, as long as they attended exactly three sessions per week.

### 2.5 Training sessions

Each training session (Figure 1D) lasted 25 minutes and was exclusively performed using the left non-dominant hand. Prior to each training session, we displayed a bar graph that illustrated the average error for each distance across their previously completed sessions. At the outset of each session, we conducted a familiarization phase where participants performed five trials of each distance. After that the LCI and HCI groups completed 270 trials during each training session, which were divided into 18 blocks of 15 trials each. During the training phase, the LCI group trained only under the blocked condition, while the HCI group trained only under the random condition.

Participants in the LCI group completed six consecutive blocks of trials with the same distances (e.g., smaller distances), followed by six consecutive blocks for each of the two remaining distances (e.g., mid and longer distances). The order of the distances was counterbalanced across participants and sessions. Participants in the HCI group trained with the three distances mixed in each of the 18 blocks (i.e., 5 trials per distance per block). Inside each block, the distances were presented in a random order.

Therefore, both groups trained each distance equally during the task; the only difference was the order in which the distances were practiced (i.e., random or blocked).

### 2.6 Test sessions

In the test session (Figure 1E), also exclusively performed using the left non-dominant hand, the task took approximately 15 minutes to be completed. A bar graph was presented to participants at the beginning of T2 displaying the average error for each distance across their previously completed sessions. At the outset of each test session, we conducted a familiarization phase where participants performed six trials of each distance. Each session comprised 108 trials, organized into 9 blocks of 12 trials. Each block included 12 trials split into 4 trials for each distance, presented in a random order.

### 2.7 Metric of performance and Behavioral statistical analysis plan

Error was quantified using absolute error, defined as the average absolute distance of the endpoint clicks relative to the target, irrespective of the direction of deviation. Lower absolute error denotes higher accuracy (Young et al., 1993). We excluded trials initiated before the target cross turned green and those with an excessive response delay (i.e., exceeding 20% of the response time computed in the familiarization trials). To investigate performance changes between T1 and T2 and among groups, we used linear mixed models (LMM) for each condition. Specifically, we modeled LMMs for absolute error, with group (HCI, LCI) and session (T1, T2) as fixed factors, and distance (short, mid, long) as well as subjects as a random factor (random intercept). We tested the main and interaction effects on the response variables, and in the case of a significant interaction, we computed pairwise comparisons and applied Holm-Bonferroni corrections to control for Type I errors. The statistical analyses were performed using Jamovi 2.3.18 (The jamovi project, 2023).

### 2.8 EEG recording and preprocessing

To ensure interpretability, we confirmed through a *t*-test that reaction times were not significantly different between the HCI and LCI groups at T1 (Δ = −0.021 ms, *t*(103) = −1.88, *p* = 0.063) and T2 (Δ = 0.014 ms, *t*(103) = 1.45, *p* = 0.150). Matching reaction times between groups in EEG studies is crucial because it ensures that any differences in brain activity are not due to differences in how quickly participants respond to stimuli. We recorded continuous EEG during the task at a rate of 1024 Hz with 24-bit A/D conversion using 128 electrodes (Biosemi ActiveTwo system) arranged according to the international 10–20 system. Two additional electrodes were used (active CMS: common-mode sense and passive DRL: driven right leg) to form a feedback loop for the amplifier reference. Brain Vision Analyzer (BrainVision Analyzer, Version 2.2.2, Brain Products GmbH, Gilching, Germany) was used for the preprocessing steps. The raw EEG signals were filtered between 0.5 and 45 Hz with 50 Hz notch using a zero-phase shift second-order Butterworth filter, and downsampled to 512 Hz. Eye blinks and saccades artifacts were corrected using independent component analysis (ICA) (Cardoso, 1998) and bad electrodes replaced using linear spline interpolation of adjacent electrodes (Perrin et al., 1987). The dataset was segmented into epochs of 2500 ms, spanning from 200 ms before the onset of the warning stimulus (i.e., the appearance of the red target cross) to 550 ms after the imperative stimulus (i.e., the color change of the cross from red to green). We excluded epochs containing electrical potential values exceeding ±80 μV, in addition to trials removed at the behavioral level. We then recalculated the data to obtain an average reference. After preprocessing, the mean number of remaining trials in the random condition for the LCI group was 82.44 (76.34%) ± 18.56 at T1 and 88.16 (81.64%) ± 14.05 at T2. For the HCI group, it was 89.53 (82.90%) ± 12.67 at T1 and 97.88 (90.63%) ± 9.36 at T2.

### 2.9 Topographical analyses

We computed an event-related potential (ERP) in the BrainVision software by averaging the epochs for each subject. Subsequently, we conducted ERP analyses using RAGU (Randomization Graphical User interfaces) software implemented in MATLAB (http://www.mathworks.com/) (Koenig et al., 2011; Habermann et al., 2018). To evaluate the results, we performed non-parametric randomization statistical tests, including the topographic consistency test (TCT), topographic analysis of variance (TANOVA), and global field power (GFP) analysis (Habermann et al., 2018; Koenig et al., 2011). All analyses were conducted on the 2500 ms epochs using 10’000 randomization runs, with a *p*-value threshold set at 0.05 to ensure robustness against Type I errors. Similar procedures employing RAGU have proven effective in identifying neural electrocortical correlates in different contexts (Ruggeri et al., 2019; Ruggeri et al., 2020; Di Muccio et al., 2022; Simonet et al., 2022; Raynal et al., 2024)

We conducted the TCT separately on each of the four ERPs (i.e., one per group for both sessions) to identify time intervals characterized by a consistent pattern of active sources (i.e., topographically similar scalp potential distribution) among participants. This ensured that the stimulus presentation consistently activated common neuronal sources across subjects over time, thereby minimizing the risk of drawing incorrect conclusions from TANOVA and GFP analyses (Habermann et al., 2018).

To investigate differences in the topographic distributions of brain activity in the random condition between the HCI and LCI groups at T1 and T2, we utilized TANOVA analysis structured with a 2 (group: HCI, LCI) × 2 (session: T1, T2). TANOVA analysis is a non-parametric randomization test that evaluates global dissimilarities in electric field topography without requiring the pre-selection of specific time points for event-related potential (ERP) analysis. Unlike channel-wise comparisons that identify differences at individual electrode levels, TANOVA assesses global dissimilarity across the entire electric field topography between groups (HCI vs LCI) and sessions (T1 vs T2) as well as their interaction, testing for significant topographic distinctions at each time point. This methodology enables the identification of significant differences resulting from variations in the active sources of the evoked potentials, rather than mere differences in source strength. To this end, the analysis was performed on amplitude-normalized maps (Global Field Power, GFP = 1) to ensure that observed topographical differences were not influenced by global field strength. GFP analysis quantitatively assesses the overall strength of neural activation between the “HCI” and “LCI” groups.

We conducted a GFP analysis to quantitatively assess the overall strength of neural activation and its modulation differences between the groups. The analysis followed a 2 (groups: HCI, LCI) × 2 (sessions: T1, T2) design. GFP is the standard deviation of all EEG electrode potentials at a single time point, providing a single value that represents the total electric field strength at that moment. GFP analysis quantifies overall electric field power without considering its spatial distribution. This is distinct from TANOVA, which investigates the spatial patterns of neural activation by examining the dissimilarity in electric field topographies across the scalp. Together, GFP and TANOVA provide a holistic view of neural dynamics, with GFP contributing to a measure of neural activation strength and TANOVA detailing the spatial arrangement of this activity.

Finally, we classified the dynamic changes in ERP topographies through microstate analysis, which decomposes the ERP signal into a set of stable topographical map configurations called microstates. For this analysis, we assumed that the topographies might vary between groups in their duration. We computed the microstate clustering using the algorithm implemented in RAGU. First, the algorithm identifies the optimal number of microstate prototype maps through a cross-validation procedure—an iterative technique that requires randomly splitting the available ERPs into learning and testing sets to determine their topographical patterns. We used the *k*-means algorithm with 100 random initializations to identify each microstate map.

### 2.10 Source analysis

For the periods marked by significant topographic differences we employed the weighted minimum-norm estimation (WMNE) method (Baillet et al., 2001) for neural source estimation, using Brainstorm software (Tadel et al., 2011). This approach offers a stable and conservative inverse solution that is widely applicable across various studies (Baillet et al., 2001). Given the absence of individual MRI templates, we constructed a generic head model using the OpenMEEG Boundary Element Method (BEM) (Kybic et al., 2005; Gramfort et al., 2010). This model assumes standard conductivities for the scalp and brain layers (1.0000 S/m) and assigns a conductivity of 0.0125 S/m for the skull layer. We applied the WMNE method assuming independent noise across electrodes (noise covariance modeled with an identity matrix) to facilitate a stable inverse solution, alongside fixed source orientation and depth weighting (exponent of 0.5 and a weight limit of 10) to accurately localize neural activity. We set the regularization parameter at a noise level of 0.1 (Baillet et al., 2001). For the source comparison between groups, at each session, in each of the significant periods, we conducted statistical comparisons using independent parametric tests, with a significance level set at *p*<0.05 and adjustments for multiple comparisons through the False Discovery Rate (FDR) method (Tadel et al., 2019).

### 2.11 Connectivity Analysis

To analyze the connectivity differences between the HCI and LCI groups, we initially calculated phase-locking value (PLV) matrices using Brainstorm software (Tadel et al., 2011). This involved transforming EEG data into source space using the Desikan-Killiany atlas (Desikan et al., 2006), composed of 68 cortical regions. Such delineation facilitates a detailed analysis of connectivity patterns across the cortex by evaluating the synchronization among these regions, represented in a 68×68 connectivity matrix. PLVs were computed within specific frequency bands after decomposing the EEG signals using the Hilbert transform. This approach enabled us to assess phase synchronization in the theta (4-7 Hz), alpha (8-12 Hz), beta (13-30 Hz), and gamma (30-45 Hz) frequency bands. Due to the long duration of the theta and alpha cycles, as in Cretton et al. (2024 submitted), we conservatively computed the theta and alpha PLVs during the whole and half epochs. Because their cycles are shorter, for the beta and gamma bands, we computed the PLVs within the time window identified as significant by TANOVA and GFP. The PLVs were calculated for each trial within the selected time. We then concatenated these PLVs to derive a single PLV value per subject, for each session, in each frequency band. To examine connectivity variations between the HCI and LCI groups, we conducted directional paired t-tests using Network-Based Statistic (NBS) software (NBS toolbox, https://www.nitrc.org/projects/nbs/) (Zalesky et al., 2010) on PLVs across selected time windows and frequency bands. The analysis was conducted in both directions—HCI > LCI and LCI > HCI—for each session (T1, T2) and frequency band. A significance level of *p* < 0.05 was set, employing 10,000 permutations, and controlling for multiple comparisons with the family-wise error rate (FWER) to reduce the probability of false network identification.

## 2. Results

### 3.1 Behavioral results

For the mixed model we found no significant effect of group on Error (*F*(1, 33.0) = 0.482, *p* = 0.492). However, there was a significant main effect of session (*F*(1, 171.0) = 103.051, *p* < .001) and a significant interaction between group and session (*F*(1, 171.0) = 8.071, *p* = 0.005). The post-hoc t-test on the fixed factor session indicated that Error decreased from T1 to T2 (Δ = 0.687, *t*(198) = 7.16, *p* < .001). The post-hoc t-test for the interaction between group and session showed that the Error between the HCI and LCI groups at T1 (Δ = 0.0666, *t*(198) = 0.490, *p* = 0.624) was not statistically different, but it was at T2 (Δ = −0.318, *t*(198) = −2.342, *p* = 0.040). Moreover, Error decreased more from T1 to T2 in the HCI group (Δ = 0.880, *t*(198) = 6.387, *p* < .001) compared to the LCI group (Δ = 0.495, *t*(198) = 3.698, *p* < .001). Importantly, the present manuscript reports only behavioral results from the test sessions (T1 and T2), in which all participants performed under the same random condition. Full behavioral analyses across the nine training sessions and transfer tasks, are reported in detail in Cretton et al., 2025.

### 3.2 Topographical analyses

Using the TCT analyses (Figure S1), we first identified periods of consistent scalp topographies within each of the four conditions separately (HCI-T1, HCI-T2, LCI-T1, LCI-T2). We then determined the intervals where these consistent topographical patterns overlapped across all four conditions. This procedure revealed common intervals characterized by stable and consistent neural topographies across all subjects and conditions, spanning from 126 ms to 726 ms and from 1870 ms to 2250 ms. These intervals, reflecting consistent patterns across conditions, were subsequently used to constrain further analyses. The TANOVA analysis revealed a significant session x group interaction between 232 and 255 ms and between 296 and 333 ms (Figure 3A, intervals highlighted in yellow). Furthermore, the GFP analysis revealed significant session x group interactions during two distinct periods: 275 to 296 ms and 366 to 446 ms (Figure 3B, intervals highlighted in purple). For subsequent analyses, we focused exclusively on the period showing consistent topographies across subjects, as identified by the TCT, and where we observed a significant session x group interaction. Therefore, we concentrated on the 126 to 726 ms interval.

**Figure 3.**
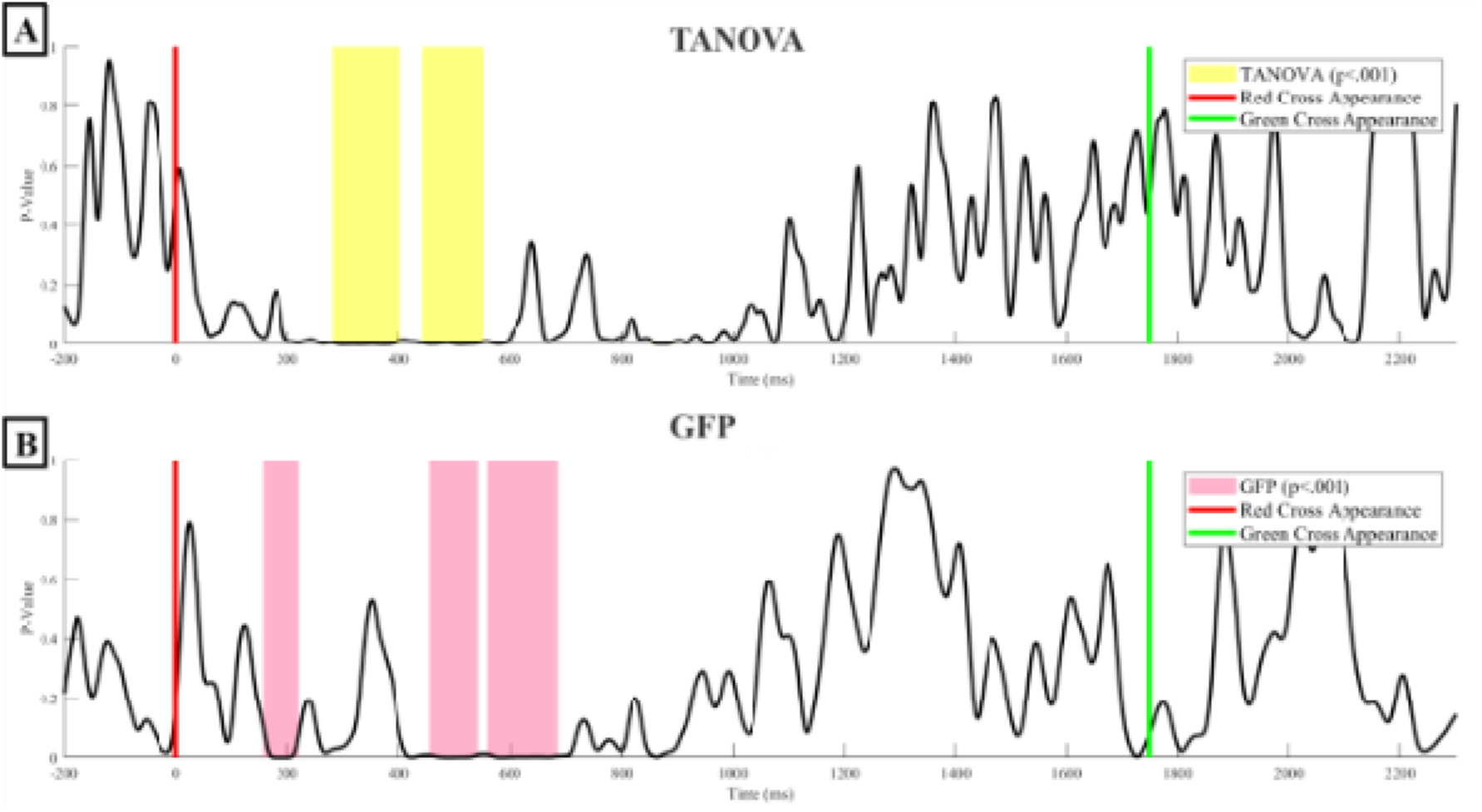
Results of TANOVA (A) and GFP (B) analyses, displaying the p-values of the differences between conditions over time. Yellow shaded areas indicate periods where TANOVA’s p-value falls below p<.05. Pink shaded areas indicate periods where GFP’s p-value falls below p<.05. The red line indicates the moment when the distance is displayed, and the green line represents the imperative stimulus signaling participants to aim at the target.

During this period, we performed microstate analysis on both groups at T1 and T2. Through cross-validation, we determined an optimal set of four microstate maps. As illustrated in Figure 4, Map 1 resembles the N1 component, characterized by posterior negativity and fronto-central positivity. Map 2 resembles the P3a component with its fronto-central positivity. Map 3 is a hybrid of the P3a-like Map 2 and the Map 4 which resembles p3b component with its centro-posterior positivity. The sequence of maps is consistent across both groups and sessions, except for the HCI group at T2, which skips Map 3 and transitions directly from Map 2 to Map 4.

**Figure 4.**
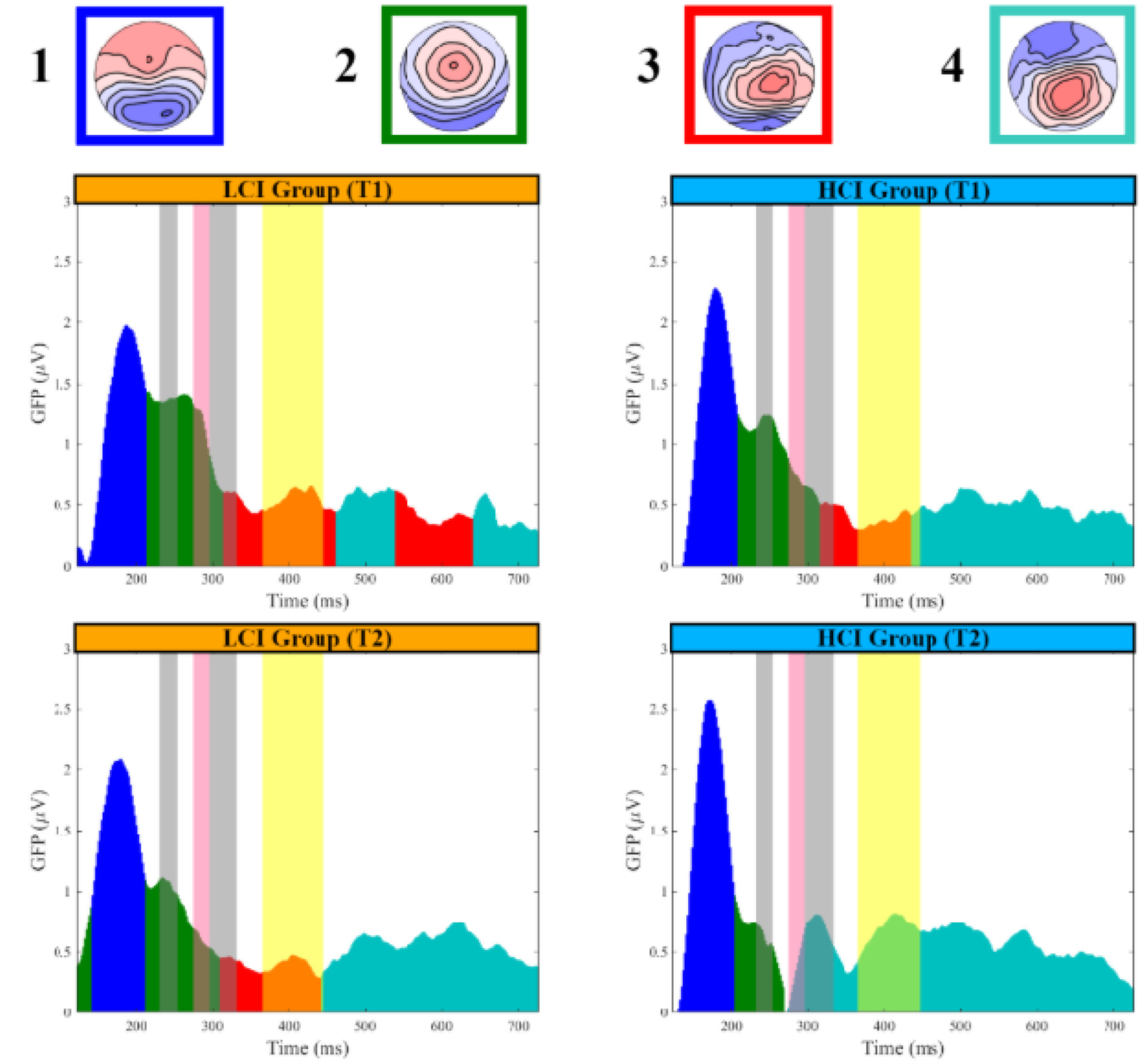
Microstates analysis (0-726ms). The four numbered microstate maps are represented as follows: Map 1 (blue), Map 2 (green), Map 3 (red), Map 4 (turquoise). The figure includes four graphs showing GFP in microvolts over time (milliseconds) for the LCI and HCI groups at T1 and T2. GFP curves are color-coded to match the microstate maps. Yellow-shaded areas highlight significant time periods (p<0.05) identified by TANOVA analysis. Pink shaded areas highlight significant time periods (p<0.05) identified by GFP analysis. The zero point corresponds to the presentation of the imperative stimulus (red cross) as shown in Figure 1.

During the N1-like topography (Map 1), no significant group and session interaction was found with neither the TANOVA or GFP analyses (Figure 4). However, during the P3a-like topography (Map 2), significant group x session interactions were detected with the TANOVA. These differences are probably driven by shorter P3a-like activity in the HCI group at T2. Specifically, Map 2 ended sooner and was shorter for the HCI group at T2 (offset at 271.1 ms, duration 66.5 ms) compared to the LCI group (offset at 310.2 ms, duration 96.2 ms). At T1, the HCI group had an offset and duration of 316 ms and 107.5 ms, respectively, while the LCI group had 314.1 ms and 99.7 ms, respectively. After Map 2, the HCI group at T2 transitioned directly to P3b-like topography (Map 4). In contrast, the HCI group at T1 and the LCI group at both T1 and T2 exhibited a transitional phase characterized by a mixture of P3a and P3b activity (Map 3). Therefore, at T2, Map 4 appear sooner in the HCI group (onset at 273 ms) compared to the LCI group (onset at 445 ms). At T1, Map 4 start at 437.2 ms in the HCI group and at 464.6 ms in the LCI group. Finally, the differences in GFP after Map 2 are likely because the HCI group at T2 had a strong P3b-like topography (Map 4), while the HCI group at T1 and the LCI group at both T1 and T2 showed a mix of weaker P3b and P3a activity (Map 3). It is why the analyses identified differences of strength of P3b activity rather than topographical differences.

The detailed dynamic representation of the topographies supports previous analyses (Figure 5). At T1, both groups showed similar topographies. However, at T2, several differences emerged. From 223 to 342 ms—a period marked by significant differences in topographies and power—the P3a-like fronto-central positivity decreased more rapidly in the HCI group than in the LCI group. During this time, the HCI group quickly transitioned to a P3b-like centro-posterior positivity, whereas the LCI group transitioned more slowly, maintaining a mix of P3a-like and P3b-like topographies. From 367 to 463 ms, a period marked by significant power differences, the HCI group exhibited a stronger P3b-like topography. In contrast, the LCI group displayed a weaker P3b-like pattern, still mixed with P3a-like characteristics. From 463 to 727 ms, both groups exhibited topographies similar to the P3b. The topography analyses are supported by single-channel ERPs at the Cz, Pz, and Fz electrodes (Figure S2). At T2, the ERP curves between groups are similar until about 200 ms, diverge between 200 ms and 500 ms, and then converge again until the end of the period.

**Figure 5.**
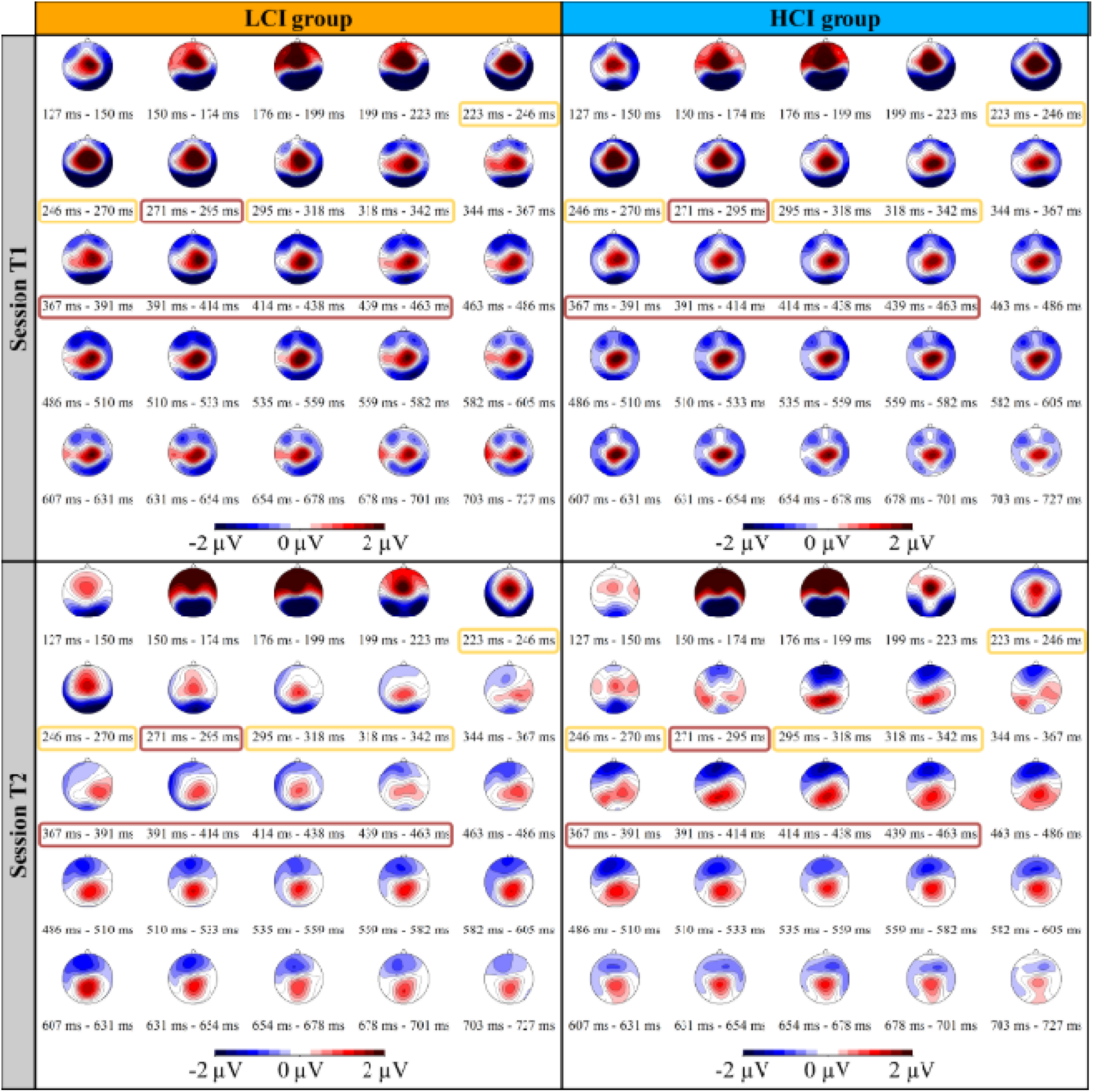
Time-course description of the topographies for LCI group at T1 (Top-left) and T2 (Bottom-left) and for HCI group at T1 (Top-right) and T2 (Bottom-right). The color-coded map illustrates the electrical scalp field potentials, in microvolts, with red indicating positive electrical potential areas and blue indicating negative electrical potential areas. Yellow squares highlight significant time periods where the p-value is below 0.05 identified with the TANOVA analysis. Rose squares highlight significant time periods where the p-value is below 0.05 identified with the GFP analysis.

We computed the average topographies for the LCI and HCI groups at both T1 and T2 for the 232 to 333 ms window (Figure 6A) and the 366 to 446 ms window (Figure 6B). Additionally, we computed standard *t*-maps to contrast these average topographies. During the 232 to 333 ms window at T1 (Figure 6A), both groups displayed a typical P3a-like topography, with no clear differences in the activity patterns. At T2, both groups showed decreased fronto-central P3a-like positivity compared to T1, but the HCI group decreased more significantly, displaying a centro-posterior P3b-like topography whereas the LCI group continued to display a P3a-like topography confirming the topographical difference found with the TANOVA.

**Figure 6.**
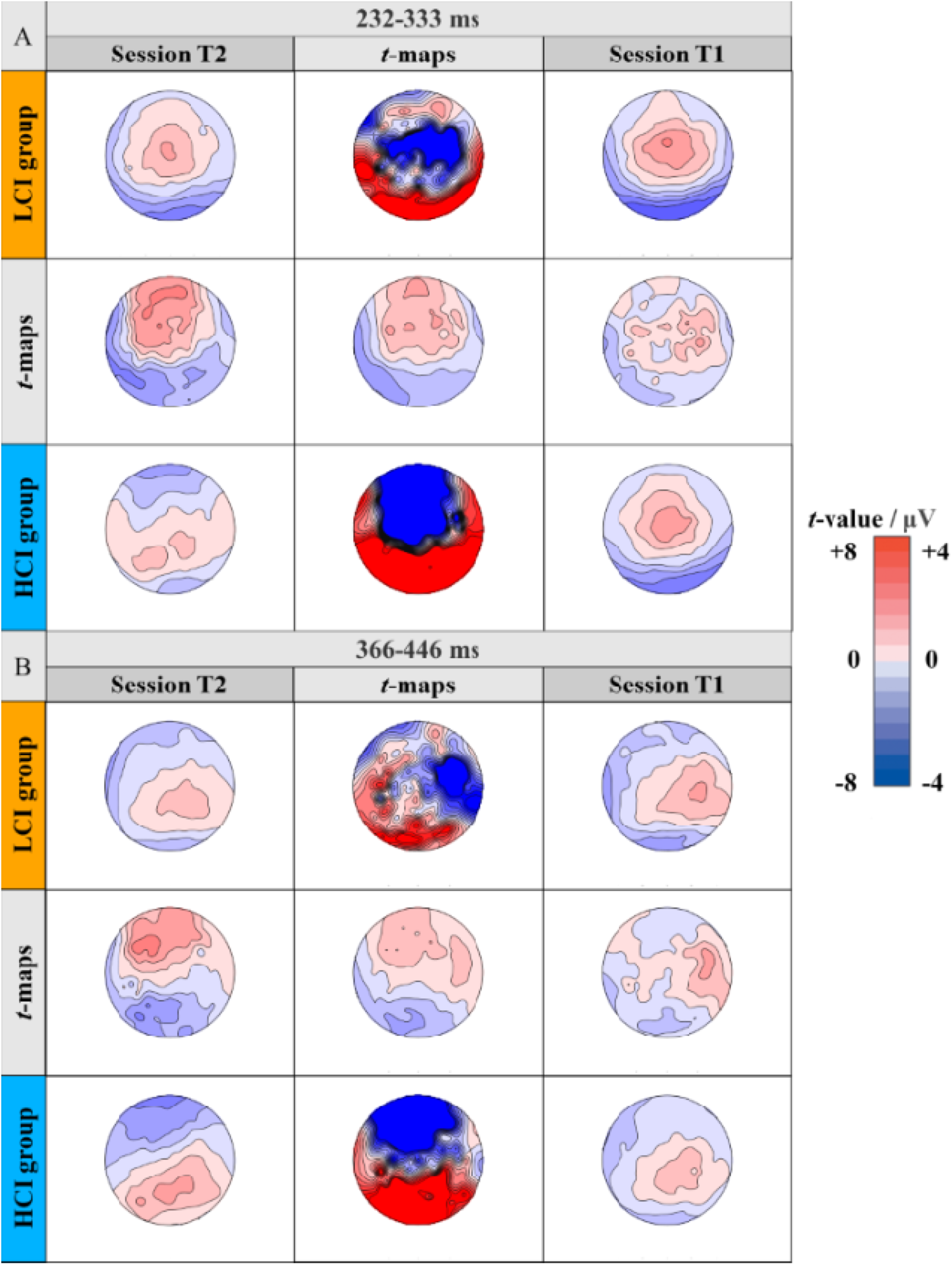
Average topographies and *t*-maps. (A) 232-333 ms period and (B) 366-446 ms period. The topographies for session T2 are displayed in the first column, while the topographies for session T1 are shown in the third column. The first row presents the topographies for the LCI group, while the third row presents the topographies for the HCI group. The middle columns and rows display *t*-maps comparing the average maps, contrasting T2 against T1 and LCI against HCI. For the average topographies, the color-coded map illustrates the electrical scalp field potentials, in microvolts, with red indicating positive electrical potential areas and blue indicating negative electrical potential areas. For the t-maps, positive (red) and negative (blue) t-values indicate respectively more positive and negative potentials in the T2 session compared to the T1 session or in the LCI group compared to the HCI group.

During the 366 to 446 ms period at T1 (Figure 6B), both groups displayed a centro-posterior P3b-like topography, likely mixed with P3a-like activity, with no clear differences between the groups. At T2, the LCI group showed no clear changes in activity patterns compared to T1. In contrast, the HCI group increased its centro-posterior activity, displaying a purer P3b-like topography. The *t*-map at T2 confirmed this, showing greater frontal activity for the LCI group and greater posterior activity for the HCI group. Both groups displayed centro-posterior P3b-like positivity, explaining why only a GFP difference, and not a topographical difference, was found during this period.

### 3.3 Source estimation analyses

As shown in Figure 7, the source estimation distribution dynamic was similar between groups at T1. This was confirmed by contrast analysis, which found no significant differences between the groups at T1 during the 232 to 332 ms or the 366 to 445 ms periods (Figure 8).

**Figure 7.**
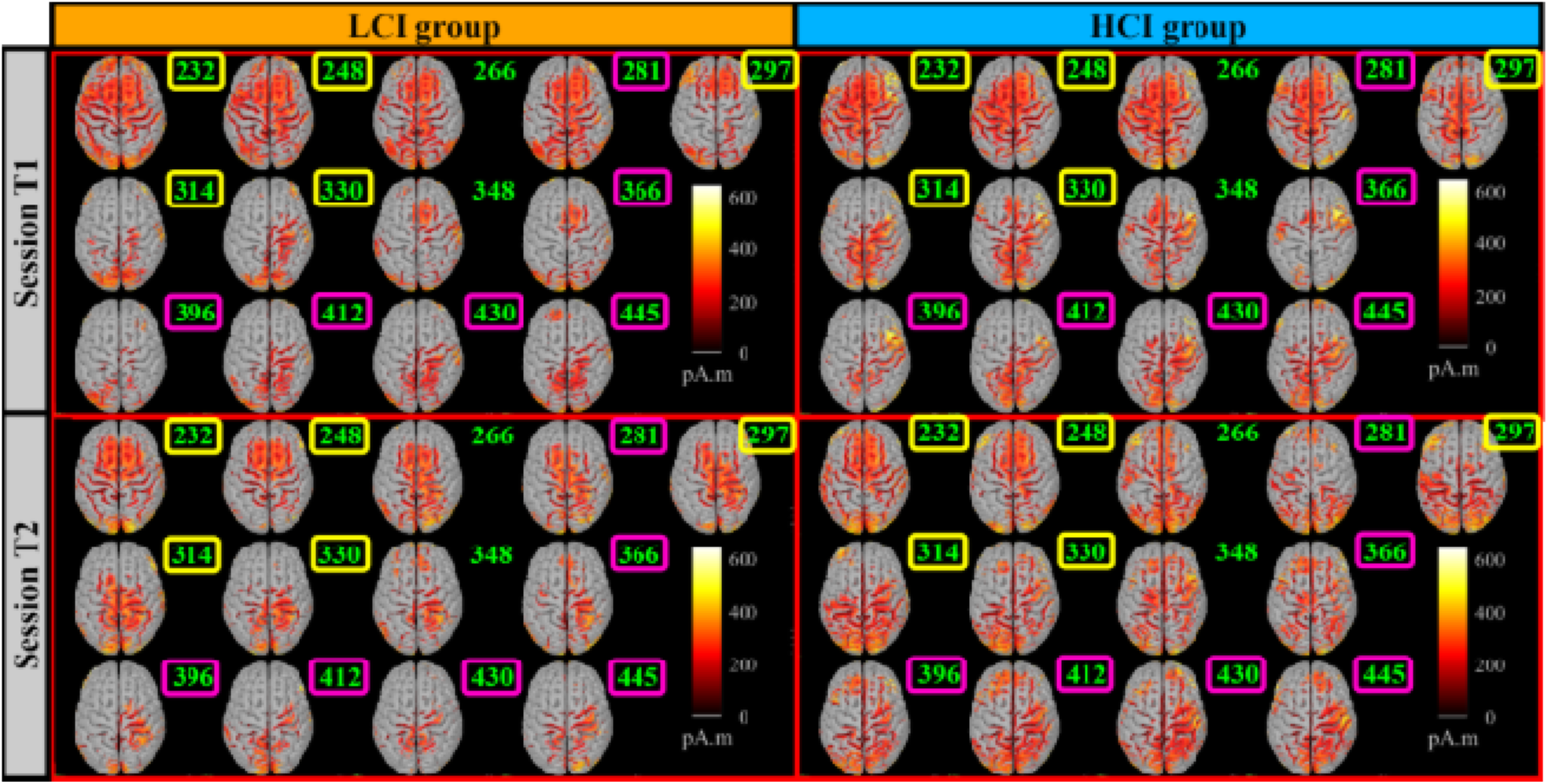
Detailed time-course of the dynamic source estimations for LCI group at T1 (Top-left) and T2 (Bottom-left) and for HCI group at T1 (Top-right) and T2 (Bottom-right). These estimations were computed using a t-test against zero at *p*<0.05, with FDR correction. The color-coded map illustrates the direction and magnitude of neural activity different from zero, measured in picoamperes times meters (pA·m). Note that this representation does not indicate the directionality of the activity (i.e., whether the potential is positive or negative). Yellow squares highlight significant time periods where the p-value is below 0.05 identified with the TANOVA analysis. Rose squares highlight significant time periods where the p-value is below 0.05 identified with the GFP analysis.

**Figure 8.**
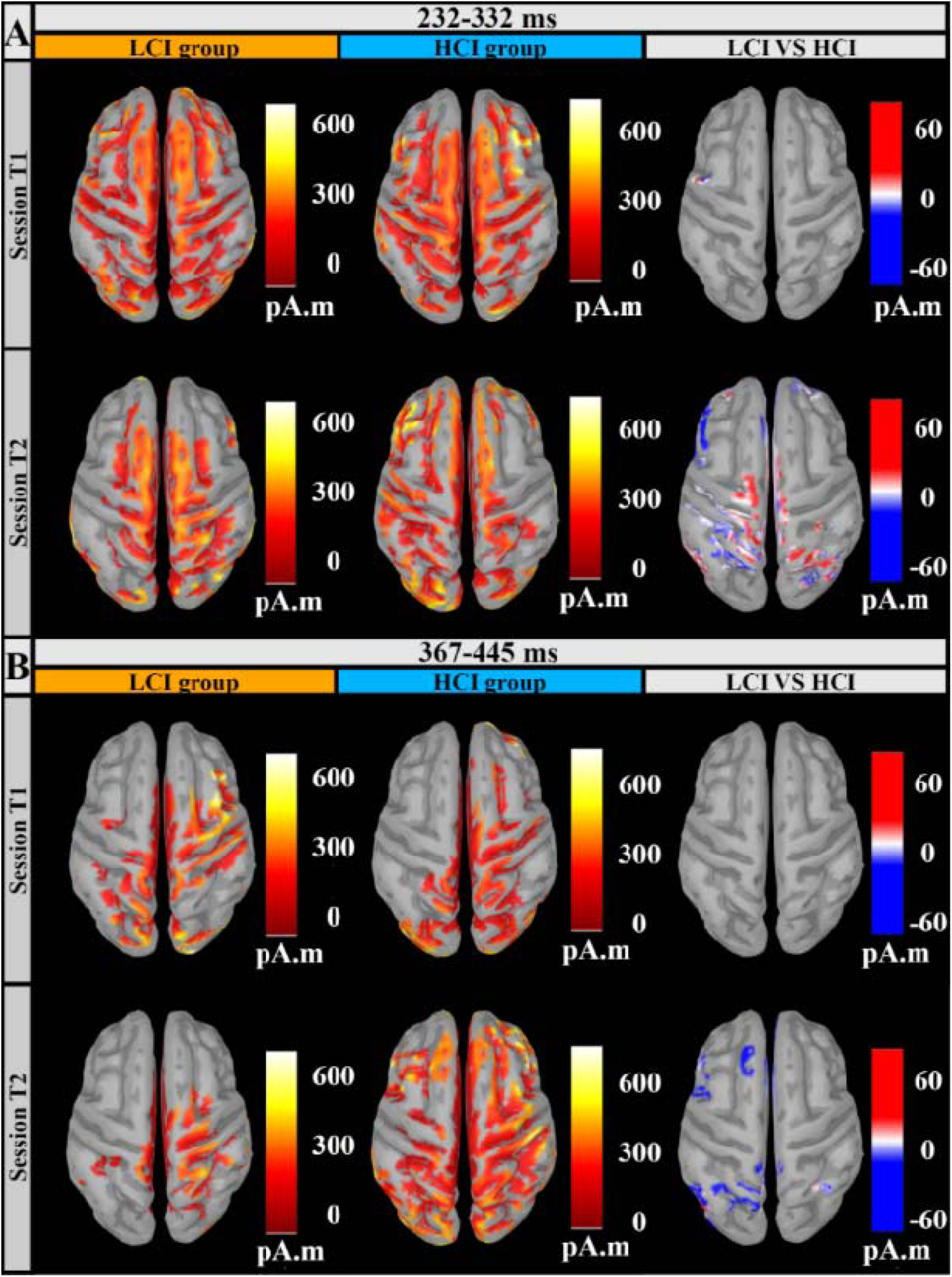
Average source estimation and contrast between groups. (A) 232-333 ms Period and (B) 366-446 ms Period. The first two columns show the source estimation for the LCI group (first column) and the HCI group (second column) at T1 (first row) and T2 (second row), computed after parametric statistics (t-test, p<0.05, with FDR correction). The color-coded map (red-yellow) illustrates the magnitude of neural activity different from zero, measured in picoamperes times meters (pA.m). The third column shows the contrast between LCI and HCI groups at T1 (first row) and T2 (second row), computed after parametric statistics (t-test, p<0.05, with FDR correction). The color-coded map indicates the direction and magnitude of differences in neural activity, measured in picoamperes times meters (pA.m). Red represents areas where there was a significantly greater electrical activity (higher pA·m values) in the LCI group than in the HCI group. Conversely, blue denotes areas showing significantly greater electrical activity in the HCI group relative to the LCI group.

However, at T2, the HCI group exhibited a faster transition from a fronto-central P3a-like activity pattern to a P3b-like centro-posterior pattern from 232 to 330 ms compared to the LCI group, as illustrated in Figure 7. Contrast analysis at T2 for the 232 to 332 ms interval showed significant differences, with the HCI group displaying more frontoparietal activation and the LCI group showing more central activation (Figure 8).

From 366 to 445 ms at T2, both groups exhibited a P3b-like centro-posterior activity pattern, but it was more pronounced in the HCI group. In addition to this greater posterior activity, the HCI group demonstrated frontal activity not observed in the LCI group, as confirmed by contrast analysis (Figure 8B).

### 3.4 Connectivity analyses

We calculated the PLVs in the theta band during the whole epoch, in the alpha band during both halves of the epoch, in the beta band from 232 to 446 ms, and in the gamma band separately between 232 and 333 ms and between 366 and 446 ms. After conducting NBS analyses using paired unidirectional t-tests across each time window, frequency band, and group at both T1 and T2, and for each direction, only one significant network was identified in the 232-333 ms time window (*p* = 0.026) (Figure 9). This network exhibited greater synchronization in the LCI group compared to the HCI group in the gamma band, involving 6 edges linking 7 nodes within an occipito-temporal-frontal network.

**Figure 9.**
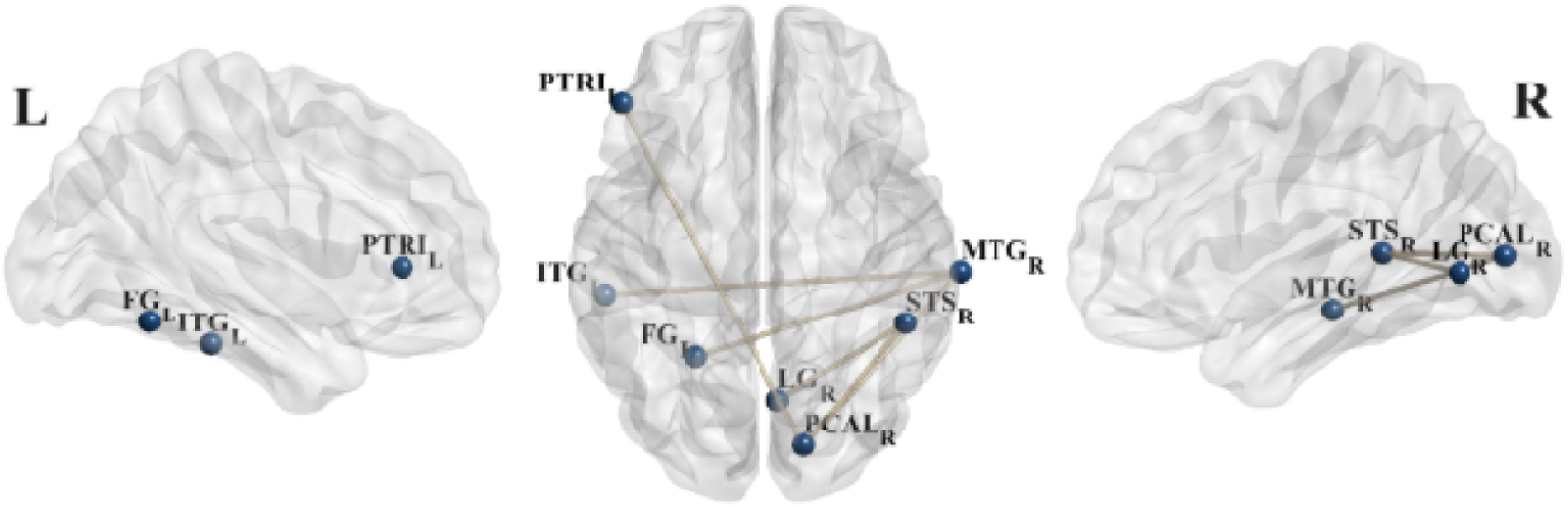
Gamma Band Network Connectivity (232-333 ms): LCI vs. HCI group at T2. This figure shows the significant network of nodes linked by edges identified with unidirectional paired t-test comparing (LCI > HCI at T2) within the gamma frequency band from 232 ms to 333 ms. R and L refer to the right and left sides, respectively. PTRl: left pars triangularis, ITGl: left inferior temporal gyrus, FGl: left fusiform gyrus, LGr: right lingual gyrus, PCALr: right pericalcarine cortex, STSr: right superior temporal sulcus, MTGr: right middle temporal gyrus.

## 3. Discussion

The aim of our study was to investigate the effects of a three-week CI training of an aiming task on neural markers of attention and working memory using a multi-scale analysis approach, including topographical, source estimation, and source connectivity analyses. We hypothesized that after training, in the random condition, the LCI group would show increased neural markers of attentional demand (N1, P3a) and working memory engagement (P3b) compared to the HCI group. Additionally, we expected that the LCI group at T2 would display a greater phase synchronization in the ventral alpha and frontoparietal theta networks throughout the trial.

Our hypotheses were partially confirmed for the differential modulation of perceptual and attentional processes as effect of CI in the random condition, but not for working memory processes. Specifically, using the TCT, we identified a period from 126 ms to 726 ms during which topographies were consistent across subjects in both groups at T1 and T2. Microstate analyses within this period revealed a succession of four stable microstates: Map 1, characterized by posterior negativity, indicative of the N1 component (Vogel & Luck, 2000; Wascher et al., 2009); Map 2, showing fronto-central positivity, characteristic of the P3a component (Halgren et al., 1998; Kok, 2001); Map 3, displaying centro-parietal positivity, a hybrid between the P3a-like Map 2 and the P3b-like Map 4, characterized by enhanced parietal positivity (Polich, 2003; Verleger, 2020). The succession of maps was consistent for the LCI group at T1 and T2 and the HCI group at T1. However, Map 3 was absent in the HCI group at T2.

Contrary to our hypotheses, no significant group x session differences in topographies or GFP were found during the N1 period. However, during the P3a-like period, significant differences were observed in both topographies and GFP. The interaction effect appears to be driven by reduced and shorter P3a-like activity in the HCI group at T2, as indicated by reduced GFP compared to the HCI group at T1 and the LCI group at both T1 and T2. Additionally, topographical and source estimation analyses showed a faster transition from P3a-like frontocentral positivity to P3b-like centroparietal positivity in the HCI group at T2. This is further supported by the absence of a transitional topography in the HCI group at T2. In contrast, at T1 and in the LCI group at both T1 and T2, this hybrid map facilitated a progressive transition from P3a-like to P3b-like topography. The GFP differences during this period likely indicate that the HCI group at T2 was already exhibiting a strong P3b, whereas at T1 and in the LCI group, this P3b component emerged more gradually.

We interpret the shorter P3a in the HCI group at T2 as indicative of a reduced attentional need for distance discrimination compared to the LCI group. The P3a component is typically observed in three-stimulus oddball tasks, where participants respond to infrequent target stimuli, frequent non-target stimuli, and infrequent novel distractor stimuli that differ from the regular stimuli. The P3a is primarily elicited by infrequent distractor stimuli, reflecting the brain’s involuntary shift in attentional allocation to assess their significance (Friedman et al., 2001; Polich & Comerchero, 2003; Polich & Criado, 2006; Polich, 2007). The amplitude of P3a increases when the discrimination between targets and non-targets is more difficult (Comerchero & Polich, 1998; Hagen et al., 2006; Sawaki & Katayama, 2009). For instance, in a study by Hagen et al. (2006), participants were presented with three types of visual stimuli: target stimuli (blue circles, 3.5 cm in diameter), standard stimuli (blue circles of varying diameters—2.7 cm for easy, 3.0 cm for medium, and 3.3 cm for hard tasks), and distractor stimuli (squares). Target and distractor stimuli had an occurrence probability of 0.12 each, while the standard stimuli appeared most frequently at a probability of 0.76. Participants were required to respond to the target stimuli by pressing a mouse key as quickly as possible and to ignore the standard and distractor stimuli. The results showed that P3a amplitude increased with greater size similarity between the target and non-targets, indicating that discrimination was more difficult. Moreover, the latency of the P3a has been shown to be greater and earlier for non-targets compared to targets (Comerchero & Polich, 1998) and to increase when performing actively compared to passively in the three-stimulus oddball task (Wronka et al., 2008). Additionally, young adults exhibit a more posterior P3 response for target stimuli and a more frontal P3 response for novel stimuli. However, with repeated trials, as in the HCI group in our study, this P3 response for targets shifts from a frontal to a more posterior scalp topography (Friedman et al., 1993; Friedman & Simpson, 1994; Fabiani & Friedman, 1995).

Although we did not find significant differences in theta or alpha networks between groups, contrary to our hypotheses, the interpretation of a reduced discrimination process in the HCI group at T2 is supported by the enhanced gamma-band synchronization in the occipito-temporo-frontal network observed during the 232 to 333 ms period in the LCI group compared to the HCI group. Gamma-band responses (GBRs) include induced components, which are not phase-locked to stimulus onset and are typically identified through single-trial analyses (for a review, see Tallon-Baudry & Bertrand, 1999; Herrmann et al., 2010). Our analysis focused specifically on these induced GBRs. The greater induced gamma-band synchronization observed in the LCI group suggests enhanced involvement of neural mechanisms supporting higher-order visual and attentional processes, consistent with prior work linking induced GBRs to stimulus discrimination and integrative visual functions (Tallon-Baudry et al., 1996, 1997; Lachaux et al., 2000; Fröunberger et al., 2007; Kim et al., 2008; Edden et al., 2009).

For instance, in a study by Tallon-Baudry et al. (1996), subjects performed a visual task involving three types of stimuli: illusory triangles, real triangles, and no-triangle stimuli. The goal was to silently count the occurrences of an additional target distractor, a curved illusory triangle, which was randomly intermixed with the three main stimuli. The study found that gamma activity occurred around 280 ms in response to both illusory and real triangles compared to the no-triangle stimulus. The authors suggest that this response, observed for triangles that are more similar to the distractor than the no-triangle stimulus, might represent the process of testing the match between the stimuli and the targets. Induced gamma activity after 200 ms was also observed in occipito-temporal electrodes in an intracranial electrode recording study where patients performed a visual discrimination task responding to triangles with inverted contours presented along with illusory and real triangles (Lachaux et al., 2000). Moreover, gamma frequency in the visual cortex has been shown to correlate with performance in an orientation discrimination task where participants had to judge if the second of two sequentially presented circular gratings was rotated clockwise or counterclockwise compared to the first (Edden et al., 2009). Importantly, similar to our results, Kim et al. (2008) reported greater induced posterior-frontal phase synchronization during the P3 latency (maximum between 300 to 500 ms) in a visual oddball task where subjects were asked to discriminate between standard and target circles varying in size.

Despite finding that P3b component was present in both groups at T2, we observed that the transition from P3a to P3b and the onset of P3b was faster in the HCI group due to the shorter duration of preceding processes. The P3b component typically appears in three-stimulus oddball tasks, where participants respond to infrequent target stimuli among regular non-targets. It is hypothesized to index the brain’s update of context within working memory in response to a mismatch between an expected schema and an actual task-relevant stimulus (Polich, 2003; Polich & Criado, 2006; Polich, 2007). Interestingly, the P3b component has also been suggested to reflect the remapping of target locations when those location change (Heath et al., 2012; Krigolson et al., 2015). Thus, at T2, the greater efficiency observed in the discrimination process within the HCI group may facilitate the initiation of updating the changing starting point location in the random condition. This interpretation is supported by the greater frontal and parietal activity observed in the HCI group during the same time period, regions known to be involved in spatial coordination transformations (Andersen & Buneo, 2002; Cohen & Andersen, 2002; Andersen & Cui, 2009). Furthermore, the reduced coverage of microstate 2 and disappearance of microstate 3 in the HCI group suggest that these participants relied on a well-learned and consolidated task-set, allowing them to more rapidly retrieve and update spatial parameters (e.g., target distance and starting location) within working memory. Rather than engaging in active reconfiguration of task rules, the HCI group may have accessed an already stabilized task-set, facilitating efficient adjustment to trial-by-trial changes.

Some limitations and methodological considerations should be noted when interpreting the results of this study. For instance, a potential limitation of our study concerns differences in timing intervals (fixation, movement preparation, and feedback periods) between the training and test sessions. Specifically, during training, we shortened and randomized these intervals primarily to reduce session duration and to limit anticipation responses. In contrast, intervals during test sessions remained longer and fixed to allow proper averaging of EEG signals, a methodological requirement when processing electrophysiological data. This difference in timing may influence interpretation of our results regarding the CI effect. Indeed, previous research on non-sequential tasks has shown that contextual interference primarily influences processes involved in preprogramming movements before execution (Immink & Wright, 1998; Immink & Wright, 2001; Wright et al., 2004). For example, Immink and Wright (2001) demonstrated that, in self-paced tasks, longer movement preparation times are observed under random practice conditions compared to blocked conditions, presumably due to the greater programming demands imposed by random practice. In the present study, because participants were provided longer preparation intervals during the test phase than during the training phase, our test sessions could be considered closer to transfer tests rather than pure retention tests. These longer intervals might have facilitated movement preprogramming, potentially reducing the differences expected between the HCI and LCI groups. Consequently, rather than accentuating between-group differences, our protocol might have unintentionally benefited the LCI group by providing participants more time to preprogram their movements, thus partially mitigating performance and neural differences typically associated with the CI effect. Additionally, altering preparation intervals might have influenced the neural correlates measured by EEG. For instance, components such as N1 and P3, which are sensitive to early perceptual encoding and attentional processing respectively, could be impacted by variations in preparation times. Shorter intervals during training might have increased the pressure on preprogramming processes. Conversely, longer preparation intervals during test sessions could have reduced these demands, potentially attenuating ERP differences between groups.

Additionally, target distances were fixed across participants (7 cm, 14 cm, and 21 cm) and not scaled relative to individual arm length. Although posture was standardized across participants using a chin rest and arm support, individual differences in seated height or limb proportions may have introduced inter-individual variability in movement execution and planning demands.

Moreover, while the LCI group practiced each distance in a blocked fashion within blocks, the practiced distance changed every six blocks. As a result, although the design maintained low contextual interference within blocks, some degree of interference was introduced across blocks. This means the LCI condition did not practice a fully repetitive practice schedule across the entire training period. Such between-blocks interleaving may have partially reduced the contrast between the blocked and random conditions, potentially contributing to the absence of stronger group × session interaction effects. The decision to implement a low, rather than no, contextual interference design was intended to sustain participant engagement and motivation throughout the extended training protocol. Additionally, differences between the HCI and LCI groups observed during the random condition at T2 may have been reduced because, prior to the left-hand data collection reported here, both groups performed a transfer task with their right hand that included 108 blocked-condition and 108 random-condition trials. Consequently, the LCI group gained additional exposure to the random condition, which may have diminished the expected neural and behavioral differences between the two groups at T2. For further details about the transfer task, readers are referred to Cretton et al. (2025).

Additionally, while our multiscale EEG analysis approach provided rich information about the dynamics of neural processes at the scalp, source, and network levels, we acknowledge that our interpretation relied heavily on classical ERP terminology (e.g., N1, P3a, and P3b). Although this facilitated interpretation within established cognitive frameworks, it may have constrained our ability to fully capture the complexity and novel insights afforded by a genuinely multiscale analysis. However, this article distinctively presents simultaneous EEG analyses at different scales, which is innovative in the contextual interference literature and could serve as a reference for formulating future hypotheses within experimental paradigm where motor parameters are manipulated.

Moreover, our attempts to link microstate maps to specific ERP components remain speculative, as the correspondence between the two is not yet firmly established. This speculative attribution underscores the need for caution in drawing direct parallels between microstate dynamics and canonical ERP markers.

Another limitation concerns the restricted time windows used for analyzing frequency-band connectivity (alpha, beta, and gamma). These intervals were selected based on significant ERP-based topographic (TANOVA) and amplitude (GFP) differences. However, oscillatory neural dynamics related to contextual interference may extend beyond these periods, and our constrained approach may have overlooked relevant differences occurring earlier or later in the trial. Future studies should therefore consider analyzing frequency-band connectivity across the entire epoch to fully characterize the temporal extent and dynamics of neural synchronization patterns associated with motor learning under different practice conditions.

A further limitation of our study arises from the fact that our reported EEG analyses focused exclusively on the random condition during the test sessions (T1 and T2). Although participants in both groups performed both random and blocked conditions during these sessions, our selective analysis might mean that observed group differences primarily reflect task-specific familiarity and practice effects associated with the random condition, rather than broader context-dependent learning adaptations. Future research should therefore explicitly test whether training-induced neural modulations generalize beyond practiced conditions, such as to new distances or effectors rather than simply transfer of condition, assessing the broader transferability of contextual interference effects.

## 4. Conclusion

In conclusion, the shorter P3a latency and lower gamma-band synchronization in the occipito-temporo-frontal network observed at T2 in the HCI group, compared to the LCI group, suggest a more facilitated discrimination process. Indeed, the HCI group, during nine training sessions with constantly changing distances in the random condition, had to discriminate between them at each trial. Conversely, the LCI group, trained exclusively in the blocked condition, did not experience this need for discrimination and therefore did not train this process. These findings align with the CI literature, which shows greater brain activity during random compared to blocked conditions during training, but more facilitated and reduced brain activity during in retention for participants trained in the random condition (Lin et al., 2011; Lin et al., 2013; Frömer et al., 2016; Lin et al., 2016; Shewokis et al., 2017; Chalavi et al., 2018; Immink et al., 2021). One limitation of our study is the potential contamination of the EEG data during the 24-hour retention session due to participants in the LCI group practicing 108 trials in the random condition during baseline. However, we believe this contamination effect is negligible when compared to the 2,538 trials completed by the HCI group in the random condition, spread across nine training sessions over three weeks, with 21 days for offline consolidation. Furthermore, we view it as a strength of our protocol that the LCI group had already experienced 108 trials in the random condition during baseline. This prior exposure reduced the potential confounding effect of encountering an untrained condition during the 24-hour retention session, allowing us to more precisely target differences in learning induced by the practice schedule rather than merely adaptation to an unfamiliar task. Moreover, while our results suggest an improvement in distance discrimination in the HCI group, whether and how it contributes to the improvement of learning efficiency remains to be investigated. Finally, considering the crucial role of the cortico-cerebral-basal ganglia network in motor learning, future longitudinal studies would benefit from incorporating approaches such as EEG-MRI co-registration. This would enable the precise localization of deep brain structures—something that high-density EEG alone cannot reliably achieve at present.

## References

Andersen, R. A., & Buneo, C. A. (2002). Intentional Maps in Posterior Parietal Cortex. Annual Review of Neuroscience, 25(Volume 25, 2002), 189–220. 10.1146/annurev.neuro.25.112701.142922

Andersen, R. A., & Cui, H. (2009). Intention, action planning, and decision making in parietal-frontal circuits. Neuron, 63(5), 568–583. 10.1016/j.neuron.2009.08.028

Baker, D. H., Vilidaite, G., Lygo, F. A., Smith, A. K., Flack, T. R., Gouws, A. D., & Andrews, T. J. (2021). Power Contours: Optimising Sample Size and Precision in Experimental Psychology and Human Neuroscience. Psychological Methods, 26(3), 295–314. 10.1037/met0000337

Baillet, S., Mosher, J. C., & Leahy, R. M. (2001). Electromagnetic brain mapping. IEEE Signal Processing Magazine, 18(6), 14–30. 10.1109/79.962275

Battig, W. F. (1979). The Flexibility of Human Memory. In Levels of Processing in Human Memory (PLE: Memory). Psychology Press.

Cardoso, J.-F. (1998). Blind signal separation: Statistical principles. Proceedings of the IEEE, 86(10), 2009–2025. Proceedings of the IEEE. 10.1109/5.720250

Chalavi, S., Pauwels, L., Heise, K.-F., Zivari Adab, H., Maes, C., Puts, N. A. J., Edden, R. A. E., & Swinnen, S. P. (2018). The neurochemical basis of the contextual interference effect. Neurobiology of Aging, 66, 85–96. 10.1016/j.neurobiolaging.2018.02.014

Cohen, Y. E., & Andersen, R. A. (2002). A common reference frame for movement plans in the posterior parietal cortex. Nature Reviews Neuroscience, 3(7), Article 7. 10.1038/nrn873

Comerchero, M. D., & Polich, J. (1998). P3a, perceptual distinctiveness, and stimulus modality. Cognitive Brain Research, 7(1), 41–48. 10.1016/S0926-6410(98)00009-3

Constantinidis, C., & Klingberg, T. (2016). The neuroscience of working memory capacity and training. Nature Reviews Neuroscience, 17(7), Article 7. 10.1038/nrn.2016.43

Cretton, A., Ruggeri, P., Brandner, C., & Barral, J. (2025). When Random Practice Makes you More Skilled: Applying the Contextual Interference Principle to a Simple Aiming Task Learning. Journal of Cognitive Enhancement. 10.1007/s41465-025-00317-5

Cretton, A., Schipper, K., Hassan, M., Ruggeri, P., & Barral, J. (2024). Enhancing perceptual, attentional, and working memory demands through variable practice schedules: Insights from high-density EEG multi-scale analyses. Cerebral Cortex, 34(11), bhae425. 10.1093/cercor/bhae425

Czyż, S. H., Wójcik, A. M., Solarská, P., & Kiper, P. (2024). High contextual interference improves retention in motor learning: Systematic review and meta-analysis. Scientific Reports, 14(1), 15974. 10.1038/s41598-024-65753-3

Desikan, R. S., Ségonne, F., Fischl, B., Quinn, B. T., Dickerson, B. C., Blacker, D., Buckner, R. L., Dale, A. M., Maguire, R. P., Hyman, B. T., Albert, M. S., & Killiany, R. J. (2006). An automated labeling system for subdividing the human cerebral cortex on MRI scans into gyral based regions of interest. NeuroImage, 31(3), 968–980. 10.1016/j.neuroimage.2006.01.021

Di Muccio, F., Ruggeri, P., Brandner, C., & Barral, J. (2022). Electrocortical correlates of the association between cardiorespiratory fitness and sustained attention in young adults. Neuropsychologia, 172, 108271. 10.1016/j.neuropsychologia.2022.108271

Edden, R. A. E., Muthukumaraswamy, S. D., Freeman, T. C. A., & Singh, K. D. (2009). Orientation Discrimination Performance Is Predicted by GABA Concentration and Gamma Oscillation Frequency in Human Primary Visual Cortex. Journal of Neuroscience, 29(50), 15721–15726. 10.1523/JNEUROSCI.4426-09.2009

Fabiani, M., & Friedman, D. (1995). Changes in brain activity patterns in aging: The novelty oddball. Psychophysiology, 32(6), 579–594. 10.1111/j.1469-8986.1995.tb01234.x

Freunberger, R., Klimesch, W., Sauseng, P., Griesmayr, B., Höller, Y., Pecherstorfer, T., & Hanslmayr, S. (2007). Gamma oscillatory activity in a visual discrimination task. Brain Research Bulletin, 71(6), 593–600. 10.1016/j.brainresbull.2006.11.014

Friedman, D., Cycowicz, Y. M., & Gaeta, H. (2001). The novelty P3: An event-related brain potential (ERP) sign of the brain’s evaluation of novelty. Neuroscience and Biobehavioral Reviews, 25(4), 355–373. 10.1016/s0149-7634(01)00019-7

Friedman, D., Simpson, G., & Hamberger, M. (1993). Age-related changes in scalp topography to novel and target stimuli. Psychophysiology, 30(4), 383–396. 10.1111/j.1469-8986.1993.tb02060.x

Friedman, D., & Simpson, G. V. (1994). ERP amplitude and scalp distribution to target and novel events: Effects of temporal order in young, middle-aged and older adults. Brain Research. Cognitive Brain Research, 2(1), 49–63. 10.1016/0926-6410(94)90020-5

Frömer, R., Stürmer, B., & Sommer, W. (2016). (Don’t) Mind the effort: Effects of contextual interference on ERP indicators of motor preparation. Psychophysiology, 53(10), 1577–1586. 10.1111/psyp.12703

Gramfort, A., Papadopoulo, T., Olivi, E., & Clerc, M. (2010). OpenMEEG: Opensource software for quasistatic bioelectromagnetics. BioMedical Engineering OnLine, 9(1), 45. 10.1186/1475-925X-9-45

Habermann, M., Weusmann, D., Stein, M., & Koenig, T. (2018). A Student’s Guide to Randomization Statistics for Multichannel Event-Related Potentials Using Ragu. Frontiers in Neuroscience, 12. https://www.frontiersin.org/articles/10.3389/fnins.2018.00355

Hagen, G. F., Gatherwright, J. R., Lopez, B. A., & Polich, J. (2006). P3a from visual stimuli: Task difficulty effects. International Journal of Psychophysiology: Official Journal of the International Organization of Psychophysiology, 59(1), 8–14. 10.1016/j.ijpsycho.2005.08.003

Halgren, E., Marinkovic, K., & Chauvel, P. (1998). Generators of the late cognitive potentials in auditory and visual oddball tasks. Electroencephalography and Clinical Neurophysiology, 106(2), 156–164. 10.1016/s0013-4694(97)00119-3

Heath, M., Bell, J., Holroyd, C. B., & Krigolson, O. (2012). Electroencephalographic evidence of vector inversion in antipointing. Experimental Brain Research, 221(1), 19–26. 10.1007/s00221-012-3141-5

Herrmann, C. S., Fründ, I., & Lenz, D. (2010). Human gamma-band activity: A review on cognitive and behavioral correlates and network models. Neuroscience & Biobehavioral Reviews, 34(7), 981–992. 10.1016/j.neubiorev.2009.09.001

Immink, M. A., Pointon, M., Wright, D. L., & Marino, F. E. (2021). Prefrontal Cortex Activation During Motor Sequence Learning Under Interleaved and Repetitive Practice: A Two-Channel Near-Infrared Spectroscopy Study. Frontiers in Human Neuroscience, 15. https://www.frontiersin.org/articles/10.3389/fnhum.2021.644968

Immink, M. A., Verwey, W. B., & Wright, D. L. (2020). The Neural Basis of Cognitive Efficiency in Motor Skill Performance from Early Learning to Automatic Stages. In C. S. Nam (Ed.), Neuroergonomics: Principles and Practice (pp. 221–249). Springer International Publishing. 10.1007/978-3-030-34784-0_12

Immink, M. A., & Wright, D. L. (1998). Contextual Interference: A Response Planning Account. The Quarterly Journal of Experimental Psychology Section A, 51(4), 735–754. 10.1080/713755789

Immink, M. A., & Wright, D. L. (2001). Motor programming during practice conditions high and low in contextual interference. Journal of Experimental Psychology: Human Perception and Performance, 27(2), 423–437. 10.1037/0096-1523.27.2.423

Kantak, S. S., Sullivan, K. J., Fisher, B. E., Knowlton, B. J., & Winstein, C. J. (2010). Neural substrates of motor memory consolidation depend on practice structure. Nature Neuroscience, 13(8), Article 8. 10.1038/nn.2596

Kim, K. H., Yoon, J., Kim, J. H., & Jung, K.-Y. (2008). Changes in gamma-band power and phase synchronization with the difficulty of a visual oddball task. Brain Research, 1236, 105–112. 10.1016/j.brainres.2008.07.118

Koenig, T., Kottlow, M., Stein, M., & Melie-García, L. (2011). Ragu: A Free Tool for the Analysis of EEG and MEG Event-Related Scalp Field Data Using Global Randomization Statistics [Research article]. Computational Intelligence and Neuroscience. 10.1155/2011/938925

Kok, A. (2001). On the utility of P3 amplitude as a measure of processing capacity. Psychophysiology, 38(3), 557–577. 10.1017/s0048577201990559

Krigolson, O. E., Cheng, D., & Binsted, G. (2015). The role of visual processing in motor learning and control: Insights from electroencephalography. Vision Research, 110, 277–285. 10.1016/j.visres.2014.12.024

Kybic, J., Clerc, M., Abboud, T., Faugeras, O., Keriven, R., & Papadopoulo, T. (2005). A common formalism for the Integral formulations of the forward EEG problem. IEEE Transactions on Medical Imaging, 24, 12–28. 10.1109/TMI.2004.837363

Lachaux, J.-P., Rodriguez, E., Martinerie, J., Adam, C., Hasboun, D., & Varela, F. J. (2000). A quantitative study of gamma-band activity in human intracranial recordings triggered by visual stimuli. European Journal of Neuroscience, 12(7), 2608–2622. 10.1046/j.1460-9568.2000.00163.x

Lage, G. M., Fernandes, L. A., Apolinário-Souza, T., Nogueira, N. G. H. M., & Ferreira, B. P. (2021). Mini-Review: Practice organization beyond memory processes. Brazilian Journal of Motor Behavior, 15(5), Article 5. 10.20338/bjmb.v15i5.259

Lee, T. D., Magill, R. A., & Weeks, D. J. (1985). Influence of Practice Schedule on Testing Schema Theory Predictions in Adults. Journal of Motor Behavior, 17(3), 283–299. 10.1080/00222895.1985.10735350

Lelis-Torres, N., Ugrinowitsch, H., Apolinário-Souza, T., Benda, R. N., & Lage, G. M. (2017). Task engagement and mental workload involved in variation and repetition of a motor skill. Scientific Reports, 7(1), Article 1. 10.1038/s41598-017-15343-3

Li, Y., & Wright, D. L. (2000). An assessment of the attention demands during random- and blocked-practice schedules. The Quarterly Journal of Experimental Psychology. A, Human Experimental Psychology, 53(2), 591–606. 10.1080/713755890

Lin, C.-H. J., Chiang, M.-C., Knowlton, B. J., Iacoboni, M., Udompholkul, P., & Wu, A. D. (2013). Interleaved practice enhances skill learning and the functional connectivity of fronto-parietal networks. Human Brain Mapping, 34(7), 1542–1558. 10.1002/hbm.22009

Lin, C.-H. J., Knowlton, B. J., Wu, A. D., Iacoboni, M., Yang, H.-C., Ye, Y.-L., Liu, K.-H., & Chiang, M.-C. (2016). Benefit of interleaved practice of motor skills is associated with changes in functional brain network topology that differ between younger and older adults. Neurobiology of Aging, 42, 189–198. 10.1016/j.neurobiolaging.2016.03.010

Lin, C.-H. J., Wu, A. D., Udompholkul, P., & Knowlton, B. J. (2010). Contextual interference effects in sequence learning for young and older adults. Psychology and Aging, 25(4), 929–939. 10.1037/a0020196

Lin, C.-H. (Janice), Knowlton, B. J., Chiang, M.-C., Iacoboni, M., Udompholkul, P., & Wu, A. D. (2011). Brain–behavior correlates of optimizing learning through interleaved practice. NeuroImage, 56(3), 1758–1772. 10.1016/j.neuroimage.2011.02.066

Magill, R. A., & Hall, K. G. (1990). A review of the contextual interference effect in motor skill acquisition. Human Movement Science, 9(3), 241–289. 10.1016/0167-9457(90)90005-X

Murray, M. M., Foxe, D. M., Javitt, D. C., & Foxe, J. J. (2004). Setting Boundaries: Brain Dynamics of Modal and Amodal Illusory Shape Completion in Humans. The Journal of Neuroscience, 24(31), 6898–6903. 10.1523/JNEUROSCI.1996-04.2004

Murray, M. M., Wylie, G. R., Higgins, B. A., Javitt, D. C., Schroeder, C. E., & Foxe, J. J. (2002). The Spatiotemporal Dynamics of Illusory Contour Processing: Combined High-Density Electrical Mapping, Source Analysis, and Functional Magnetic Resonance Imaging. Journal of Neuroscience, 22(12), 5055–5073. 10.1523/JNEUROSCI.22-12-05055.2002

Pauwels, L., Chalavi, S., Gooijers, J., Maes, C., Albouy, G., Sunaert, S., & Swinnen, S. P. (2018). Challenge to Promote Change: The Neural Basis of the Contextual Interference Effect in Young and Older Adults. The Journal of Neuroscience: The Official Journal of the Society for Neuroscience, 38(13), 3333–3345. 10.1523/JNEUROSCI.2640-17.2018

Peirce, J., Gray, J. R., Simpson, S., MacAskill, M., Höchenberger, R., Sogo, H., Kastman, E., & Lindeløv, J. K. (2019). PsychoPy2: Experiments in behavior made easy. Behavior Research Methods, 51(1), 195–203. 10.3758/s13428-018-01193-y

Perrin, F., Pernier, J., Bertrand, O., Giard, M. H., & Echallier, J. F. (1987). Mapping of scalp potentials by surface spline interpolation. Electroencephalography and Clinical Neurophysiology, 66(1), 75–81. 10.1016/0013-4694(87)90141-6

Polich, J. (2003). Theoretical Overview of P3a and P3b. In J. Polich (Ed.), Detection of Change: Event-Related Potential and fMRI Findings (pp. 83–98). Springer US. 10.1007/978-1-4615-0294-4_5

Polich, J. (2007). Updating P300: An integrative theory of P3a and P3b. Clinical Neurophysiology, 118(10), 2128–2148. 10.1016/j.clinph.2007.04.019

Polich, J., & Comerchero, M. D. (2003). P3a from visual stimuli: Typicality, task, and topography. Brain Topography, 15(3), 141–152. 10.1023/a:1022637732495

Polich, J., & Criado, J. R. (2006). Neuropsychology and neuropharmacology of P3a and P3b. International Journal of Psychophysiology, 60(2), 172–185. 10.1016/j.ijpsycho.2005.12.012

Ramezanzade, H., Saemi, E., Broadbent, D. P., & Porter, J. M. (2022). An Examination of the Contextual Interference Effect and the Errorless Learning Model during Motor Learning. Journal of Motor Behavior, 54(6), 719–735. 10.1080/00222895.2022.2072265

Raynal, E., Schipper, K., Brandner, C., Ruggeri, P., & Barral, J. (2024). Electrocortical correlates of attention differentiate individual capacity in associative learning. Npj Science of Learning, 9(1), 1–11. 10.1038/s41539-024-00236-8

Riley, M. R., & Constantinidis, C. (2016). Role of Prefrontal Persistent Activity in Working Memory. Frontiers in Systems Neuroscience, 9. https://www.frontiersin.org/articles/10.3389/fnsys.2015.00181

Ruggeri, P., Meziane, H. B., Koenig, T., & Brandner, C. (2019). A fine-grained time course investigation of brain dynamics during conflict monitoring. Scientific Reports, 9(1), Article 1. 10.1038/s41598-019-40277-3

Ruggeri, P., Nguyen, N., Pegna, A. J., & Brandner, C. (2020). Interindividual differences in brain dynamics of early visual processes: Impact on score accuracy in the mental rotation task. Psychophysiology, 57(11), e13658. 10.1111/psyp.13658

Sawaki, R., & Katayama, J. (2009). Difficulty of Discrimination Modulates Attentional Capture by Regulating Attentional Focus. Journal of Cognitive Neuroscience, 21(2), 359–371. 10.1162/jocn.2008.21022

Shea, J. B., & Morgan, R. L. (1979). Contextual interference effects on the acquisition, retention, and transfer of a motor skill. Journal of Experimental Psychology: Human Learning and Memory, 5(2), 179–187. 10.1037/0278-7393.5.2.179

Shea, J. B., & Zimny, S. T. (1983). Context Effects in Memory and Learning Movement Information. In R. A. Magill (Ed.), Advances in Psychology (Vol. 12, pp. 345–366). North-Holland. 10.1016/S0166-4115(08)61998-6

Shewokis, P. A., Shariff, F. U., Liu, Y., Ayaz, H., Castellanos, A., & Lind, D. S. (2017). Acquisition, retention and transfer of simulated laparoscopic tasks using fNIR and a contextual interference paradigm. The American Journal of Surgery, 213(2), 336–345. 10.1016/j.amjsurg.2016.11.043

Simonet, M., Ruggeri, P., Sallard, E., & Barral, J. (2022). The field of expertise modulates the time course of neural processes associated with inhibitory control in a sport decision-making task. Scientific Reports, 12(1), 7657. 10.1038/s41598-022-11580-3

Soderstrom, N. C., & Bjork, R. A. (2015). Learning Versus Performance: An Integrative Review. Perspectives on Psychological Science, 10(2), 176–199. 10.1177/1745691615569000

Tadel, F., Baillet, S., Mosher, J. C., Pantazis, D., & Leahy, R. M. (2011). Brainstorm: A user-friendly application for MEG/EEG analysis. Computational Intelligence and Neuroscience, 2011, 879716. 10.1155/2011/879716

Tadel, F., Bock, E., Niso, G., Mosher, J. C., Cousineau, M., Pantazis, D., Leahy, R. M., & Baillet, S. (2019). MEG/EEG Group Analysis With Brainstorm. Frontiers in Neuroscience, 13. https://www.frontiersin.org/journals/neuroscience/articles/10.3389/fnins.2019.00076

Tallon-Baudry, C., & Bertrand, O. (1999). Oscillatory gamma activity in humans and its role in object representation. Trends in Cognitive Sciences, 3(4), 151–162. 10.1016/S1364-6613(99)01299-1

Tallon-Baudry, C., Bertrand, O., Delpuech, C., & Pernier, J. (1996). Stimulus Specificity of Phase-Locked and Non-Phase-Locked 40 Hz Visual Responses in Human. The Journal of Neuroscience, 16(13), 4240–4249. 10.1523/JNEUROSCI.16-13-04240.1996

Tallon-Baudry, C., Bertrand, O., Delpuech, C., & Pernier, J. (1997). Oscillatory γ-Band (30–70 Hz) Activity Induced by a Visual Search Task in Humans. Journal of Neuroscience, 17(2), 722–734. 10.1523/JNEUROSCI.17-02-00722.1997

The jamovi project. (2023). Jamovi (Version 2.3) [Computer Software]. Retrieved from https://www.jamovi.org

Veale, J. F. (2014). Edinburgh Handedness Inventory - Short Form: A revised version based on confirmatory factor analysis. Laterality, 19(2), 164–177. 10.1080/1357650X.2013.783045

Verleger, R. (2020). Effects of relevance and response frequency on P3b amplitudes: Review of findings and comparison of hypotheses about the process reflected by P3b. Psychophysiology, 57(7), e13542. 10.1111/psyp.13542

Vogel, E. K., & Luck, S. J. (2000). The visual N1 component as an index of a discrimination process. Psychophysiology, 37(2), 190–203. 10.1111/1469-8986.3720190

Wascher, E., Hoffmann, S., Sänger, J., & Grosjean, M. (2009). Visuo-spatial processing and the N1 component of the ERP. Psychophysiology, 46(6), 1270–1277. 10.1111/j.1469-8986.2009.00874.x

Wright, D. L., Black, C. B., Immink, M. A., Brueckner, S., & Magnuson, C. (2004). Long-Term Motor Programming Improvements Occur Via Concatenation of Movement Sequences During Random But Not During Blocked Practice. Journal of Motor Behavior, 36(1), 39–50. 10.3200/JMBR.36.1.39-50

Wright, D., Verwey, W., Buchanen, J., Chen, J., Rhee, J., & Immink, M. (2016). Consolidating behavioral and neurophysiologic findings to explain the influence of contextual interference during motor sequence learning. Psychonomic Bulletin & Review, 23(1), 1–21. 10.3758/s13423-015-0887-3

Wronka, E., Kaiser, J., & Coenen, A. M. L. (2008). The auditory P3 from passive and active three-stimulus oddball paradigm. Acta Neurobiologiae Experimentalis, 68(3), 362–372. 10.55782/ane-2008-1702

Young, D. E., Cohen, M. J., & Husak, W. S. (1993). Contextual interference and motor skill acquisition: On the processes that influence retention. Human Movement Science, 12(5), 577–600. 10.1016/0167-9457(93)90005-A

Zalesky, A., Fornito, A., & Bullmore, E. T. (2010). Network-based statistic: Identifying differences in brain networks. NeuroImage, 53(4), 1197–1207. 10.1016/j.neuroimage.2010.06.041

