## Supplementary material for "Learning-Induced Effects of Practice Schedule Variability on Stimuli Discrimination Efficiency: High-Density EEG Multi-scale Analyses of Contextual Interference Effect": Table S1

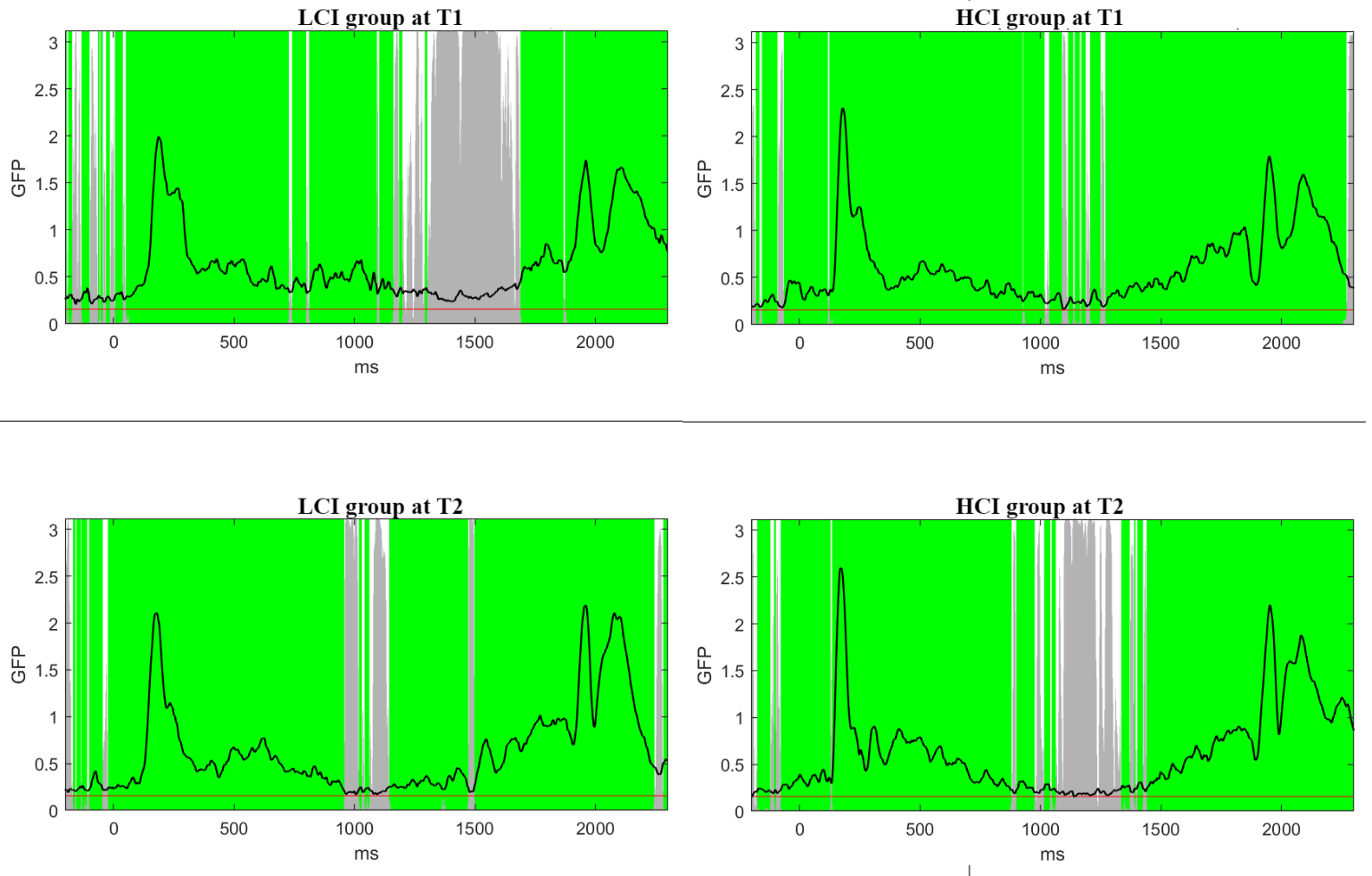

**Figure S1.** Result of Topographical Consistency Test for each group at T1 and T2. The figure depicts the GFP dynamic over time during each epoch. The green areas represent period of consistent topographies within each condition. Grey areas represents period of inconsistent topographies within each condition. The TCT analysis identified periods of consistent topographies across subjects, shared among the four ERPs, spanning 126 ms to 726 ms and 1870 ms to 2250 ms.

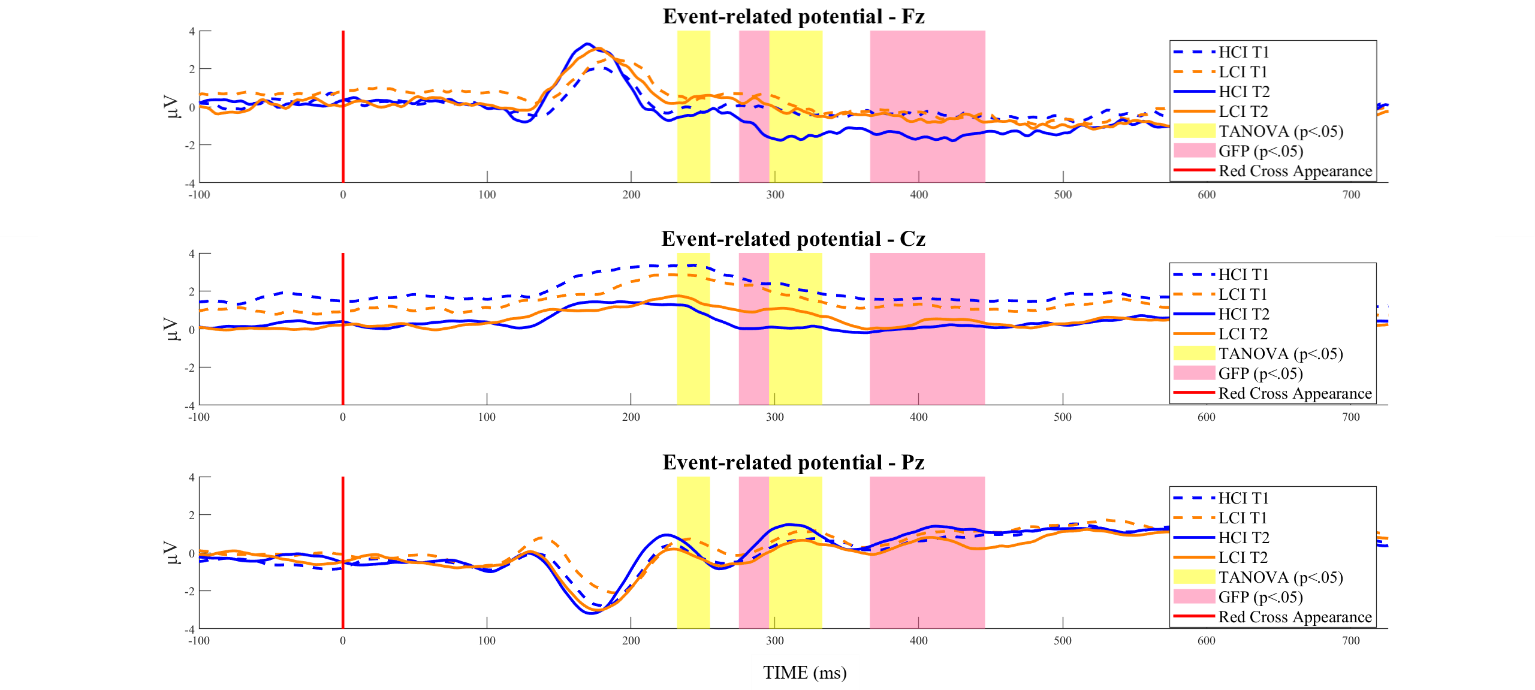

**Figure S2. Exemplar ERPs at the Cz, Fpz, and Pz electrodes for the HCI group at T1 (blue full line) and T2 (blue dashed line), and for the LCI group at T1 (orange full line) and at T2 (orange dashed line). The red line indicates when the distance is displayed, and the green line marks the imperative stimulus signaling participants to aim at the target. The time series is aligned to the onset of the imperative stimulus (0 ms). Yellow-shaded areas indicate periods where TANOVA’s *p*-value falls below *p*<.05. Rose-shaded areas indicate periods where GFP’s p-value falls below p<.05.**
